# Nitric Oxide Inhibition of Glycyl Radical Enzymes and Their Activases

**DOI:** 10.1101/2025.02.23.639758

**Authors:** Juan Carlos Cáceres, Nathan G. Michellys, Brandon L. Greene

## Abstract

Innate immune response cells produce high concentrations of the free radical nitric oxide (NO) in response to pathogen infection. The antimicrobial properties of NO include non-specific damage to essential biomolecules and specific inactivation of enzymes central to aerobic metabolism. However, the molecular targets of NO in anaerobic metabolism are less understood. Here, we demonstrate that the *Escherichia coli* glycyl radical enzyme pyruvate formate lyase (PFL), which catalyzes the anaerobic metabolism of pyruvate, is irreversibly inhibited by NO. Using electron paramagnetic resonance and site-directed mutagenesis we show that NO destroys the glycyl radical of PFL. The activation of PFL by its cognate radical S-adenosyl-L-methionine-dependent activating enzyme (PFL-AE) is also inhibited by NO, resulting in the conversion of the essential iron-sulfur cluster to dinitrosyl iron complexes. Whole-cell EPR and metabolic flux analyses of anaerobically growing *Escherichia coli* show that PFL and PFL-AE are inhibited by physiologically relevant levels of NO in bacterial cell cultures, resulting in diminished growth and a metabolic shift to lactate fermentation. The class III ribonucleotide reductase (RNR) glycyl radical enzyme and its corresponding RNR-AE are also inhibited by NO in a mechanism analogous to those observed in PFL and PFL-AE, which likely contributes to the bacteriostatic effect of NO. Based on the similarities in reactivity of the PFL/RNR and PFL-AE/RNR-AE enzymes with NO, the mechanism of inactivation by NO appears to be general to the respective enzyme classes. The results implicate an immunological role of NO in inhibiting glycyl radical enzyme chemistry in the gut.

## INTRODUCTION

Opportunistic pathogens infecting the gastrointestinal and respiratory tracks, such as *Shigella dysenteriae, Salmonella enterica, Escherichia coli, Pseudomonas aeruginosa* and *Staphylococcus aureus*, are facultative anaerobes.^1–5^ This metabolic flexibility allows for aerobic pathogen transmission and subsequent establishment and proliferation in the anaerobic or microaerobic environment of the host mucosal epithelium-microbiome interface.^6,7^ Dysregulation of the microbiome at this interface induces a host innate immune response, resulting in the release of reactive oxygen species (ROS, *e*.*g*. H_2_O_2_)^8,9^ and reactive nitrogen species (RNS, *e*.*g*. NO),^10–12^ both of which have pleiotropic and hormetic, yet still undefined roles in pathophysiology.^13,14^ For microorganisms adapted to anaerobic metabolism, ROS disrupt redox homeostasis and damage biomolecules and essential cofactors.^15^ The effect(s) of RNS, such as NO, on anaerobic metabolism are less defined and may have practical implications in therapeutic developments to manage or treat diseases. ^15–18^

During anaerobic glycolysis, many bacteria and archaea in the human gut, at least partially, metabolize pyruvate to acetyl-coenzyme A (acetyl-CoA) and formate through the action of pyruvate formate lyase (PFL).^19–21^ PFL is a member of the glycyl radical enzyme (GRE) family, requiring a post-translationally installed glycyl radical for activity.^22–24^ This essential radical is installed by a specific activating enzyme, PFL-AE, a member of the radical S-adenosyl-L-methionine (SAM)-dependent enzyme superfamily.^25–27^ The acetyl-CoA product can be used for substrate level phosphorylation to form adenosine triphosphate (ATP) and serve as an electron sink to recycle nicotinamide adenine dinucleotide (NAD^+^), or provide carbon in anabolic biosynthesis.^28^ The formate product can be expelled as a waste product, either as formate or carbon dioxide and hydrogen, or used as an electron and carbon source.^29–31^ Anaerobic growth in organisms that utilize PFL is often possible in the absence of PFL, provided a suitable electron acceptor is present, such as nitrate, or acetate is available for acetyl-CoA biosynthesis.

During infection, PFL is up-regulated in many opportunistic pathogens. The deletion of *pflA* (PFL-AE) and *pflB* (PFL) genes results in a loss of virulence in some pathogens, further suggesting PFL provides a virulent fitness advantage.^32–36^ Given its role in central metabolism and pathogenesis, and the inherent reactivity of amino acid radicals, we hypothesize that PFL is a molecular target of the immune response. The glycyl radical of PFL is extremely sensitive to O_2_, cleaving the enzyme polypeptide chain at the glycyl α-carbon, but to our knowledge, no other ROS have been investigated.^37,38^ The radical SAM PFL-AE contains an essential [Fe_4_S_4_] cofactor and is also sensitive to inactivation by O_2_, suggesting that both PFL and PFL-AE are susceptible to inactivation by O_2_, and potentially other ROS.^19^ Evidence for the reactivity of the RNS NO with PFL or PFL-AE, however, is indirect. In a study of *S. aureus*, exposure to NO under anaerobic conditions results in the diversion of metabolism from ethanol fermentation via PFL to lactate production, and in a murine infection model, *S. aureus* was non-pathogenic in the absence of an NO-inducible lactate dehydrogenase.^39^ These results suggest that NO inhibits PFL *in vivo*, with implications in pathogenic disease progression.

Here, we report the reactivity of NO with active glycyl radical-containing PFL (aPFL) from *E. coli*, PFL-AE, and additional representative members of the GRE and radical SAM enzyme families. *In vitro*, the addition of an NO donor to aPFL results in the complete loss of the glycyl radical and enzyme activity. The reaction appears to be diffusion-limited and specific to the PFL glycyl radical. We observe similar reactivity with the *E. coli* class III ribonucleotide reductase (RNR), suggesting the mechanism of irreversible inhibition is common to the GRE family. We also demonstrate that PFL-AE and RNR-AE are irreversibly inhibited by NO, although through a slower process, involving dinitrosyl iron products, known inhibition products of [Fe_4_S_4_] cluster-containing enzymes associated with aerobic metabolism such as aconitase and succinate dehydrogenase.^40–42^ The inhibition of PFL and PFL-AE was also observed *in vivo* by whole-cell electron paramagnetic resonance (EPR) spectroscopy, revealing the quenching of the glycyl radical signal and the concomitant formation of dinitrosyl iron complexes (DNICs) in PFL and PFL-AE overexpressing cells. This inhibition was accompanied by a shift in the metabolic products of anaerobically growing *E. coli* from acetate, formate, and ethanol to lactate production. These findings suggest that NO inhibits anaerobic microbial metabolism through multiple mechanisms, contributing to the antimicrobial activity of NO in the host immune response against pathogenic infections in anaerobic or microaerobic environments.

## EXPERIMENTAL

### Materials

Electrocompetent DH5α and BL21(DE3) *E. coli* and NEBuilder® HiFi DNA Assembly Master Mix were purchased from New England Biolabs. Carbenicillin and L-arabinose were purchased from GoldBio. Chloramphenicol, kanamycin and agarose were purchased from Apex Bioresearch Products. LB Miller broth, citrate synthase (porcine heart), malate dehydrogenase (porcine heart), bovine serum albumin, myoglobin (equine heart), alcohol dehydrogenase (*Saccharomyces cerevisiae*), tris(hydroxymethyl)amino-methane base (Tris), β-nicotinamide adenine dinucleotide (NAD^+^), S-(5′-adenosyl)-L-methionine (SAM) iodide salt, KH_2_PO_4_, Triton X-100, glycerol, MgCl_2_·6H_2_O, oxamic acid, sodium pyruvate, sodium formate, sodium lactate, ethanol, malic acid, iodoacetamide, LC-MS grade trifluoroacetic acid (TFA), dithiothreitol (DTT), L-cysteine, (NH_4_)Fe^II^(SO_4_)_2_, formic acid, 4-hydroxy-TEMPO, urea, sodium dithionite (NaDT), sodium borohydride (NaBH_4_), glycine, glucose, MgSO_4_, CaCl_2_, biotin, thiamin, Na_2_HPO_4_, NH_4_Cl, ethylenediaminetetraacetic acid (EDTA), FeCl_3_·6H_2_O, ZnCl_2_, CuCl_2_·2H_2_O, CoCl_2_·6H_2_O, H_3_BO_3_, MnCl_2_·6H_2_O, iron ICP standards (TraceCERT), 70% trace metal-free nitric acid, isopropyl-β-D-1-thiogalactopyranoside (IPTG), cytidine triphosphate (CTP), adenosine triphosphate (ATP), cytidine, 2′-deoxycytidine (dC), and Amicon Ultra centrifugal filter units were purchased from Millipore Sigma. 5-Deazariboflavin was obtained from Santa Cruz Biotechnologies.

Sequencing grade modified trypsin was purchased from Promega Corporation. Coenzyme A was purchased from Co-Ala Biosciences. HiTrap desalting 5 mL columns were purchased from Cytiva Life Sciences. Nickel nitrilotriacetic acid (Ni-NTA) agarose resin was purchased from Qiagen. His-Pur™ Cobalt Resin and 0.5 mL Zeba™ spin desalting columns of 7 kDa molecular weight cut-off (MWCO) were purchased from Thermo Scientific. Adenosine (A) was purchased from VWR. Diethylammonium (Z)-1-(N,N-diethylamino)diazen-1-ium-1,2-diolate (NONOate) was purchased from Cayman Chemicals. SapphireAmp® Fast PCR master mix was purchased from Takara Bio. Vivaspin 20 filtration units were purchased from Sartorius. The acetic acid assay kit (ACS Manual format) was purchased from Megazyme. Calf alkaline phosphatase was purchased from Roche. HPLC-grade water with 0.1% TFA and acetonitrile with 0.1% TFA were purchased from Honeywell. Milli-Q water (>17 MΩ) was used for preparing all solutions. The plasmid pCAL-n-EK encoding the *pflA* gene was a gift from Dr. Joan Broderick.^43^ The *pflB* gene (Uniprot ID P09373) and the mutants C_418_S and C_419_S were cloned into the plasmid pCM8 previously.^44^ The plasmid pCm2 *Nik*J was available from a previous study.^45^ The *Pseudomonas* sp. 101 formate dehydrogenase (FDH) gene (Uniprot ID P33160) was codon optimized and synthesized by Integrated DNA Technologies. The plasmids pFGET19_Ulp1 and pHYRSF53 were a gift from Hideo Iwai (Addgene plasmid # 64697 and # 64696, respectively).^46^ The TEVSH plasmid was a gift from Dr. Helena Berglund (Addgene plasmid # 125194).^47^ The plasmid pDB1282 was a gift from Dr. Squire Booker and previously constructed in the laboratory of Dr. Dennis Dean.^48,49^ The plasmid pET28a-EcNrdD and pN9-EcNrdG were a gift from Dr. JoAnne Stubbe^50^. The plasmid pLZ113 harboring the D176G-I177L-F178W LDH was a gift from Dr. Han Li.^51^

### Construction of Plasmids

To produce and purify PFL-AE, RNR-AE and FDH, we followed similar cloning strategies. Each gene was cloned into the pHYRSF53 plasmid using Gibson assembly.^52^ These plasmids yield expressed proteins with an N-terminal 6× polyhistidine tag, up-stream of a small ubiquitin-like modifying (SUMO) protein fusion, termed SUMO-PFL-AE, SUMO-RNR-AE and SUMO-FDH respectively. The genes were cloned into the plasmid replacing the gene of the protein of interest encoded in pHYRSF53, downstream of the SUMO gene and in the same open reading frame by Gibson assembly. The DNA fragment containing the genes (fragments) and the plasmid backbone (vector) were cloned using the primers listed in the **Supporting Information Table S1**.

The PCR-amplified complementary DNA fragments were assembled using NEBuilder® HiFi DNA Assembly Master Mix following the manufacturer’s instructions. Assembled reaction products were transformed into *E. coli* DH5α cells and streaked onto LB-agar plates supplemented with 50 μg/mL kanamycin. Successful transformants were identified using colony PCR with the SapphireAmp fast PCR-hot-start master mix, as directed by the manufacturer, and the fidelity of the cloning was confirmed by Sanger sequencing through UC Berkeley DNA Sequencing Facility using the T7 promoter/terminator primers.

### Protein Expression and Purification

The expression and purification of the TEV protease and the ubiquitin-like protease 1 (ULP1) were performed as previously described.^46,47^ The protein expression and purification of PFL wild type (wt, 77 ± 3 μmol/min×mg normalized to G•) and the serine (S) mutants C_418_S and C_419_S, as well as PFL-AE, were carried out as previously described without further modifications.^44^ The enzyme NikJ (2.6 ± 0.1 Fe/protein) was expressed and purified as reported before using the pCm2 plasmid.^45^ Lactate dehydrogenase (LDH, 1,330 ± 30 s^−1^) was expressed and purified as described before.^53^ Expression and purification of FDH (2.3 μmol/min×mg), RNR (1,350 ± 90 nmol/min×mg normalized to G•), RNR-AE (2.2 ± 0.1 Fe/protein) and PFL-AE (2.1 ± 0.2 Fe/protein) are detailed in **Supporting Information Supplemental Methods**. Pure proteins concentration in water was determined using the Edelhoch method^54^ but using the extinction coefficients for tryptophan (W) and tyrosine (Y) determined by Pace.^55^

### Iron Quantification

We quantified iron by inductively coupled plasma optical emission spectroscopy (ICP-OES). Typically, 8 nmol of protein was digested by adding 86 μL of 70% (v/v) trace metal-free nitric acid and incubated overnight at room temperature followed by 2 h incubation at 90 °C. Once samples returned to room temperature, 70 μL of 30% (v/v) hydrogen peroxide was added and incubated at 90 °C for 1 h. Finally, water was added to a final weight of 3 g and analyzed in an Agilent 5800 ICP-OES in the following configuration: read time 5 s, RF power 1.45 kW, stabilization time 15 s, in axial viewing mode, nebulizer flow 0.7 L/min, plasma flow 12 L/min, and auxiliary flow 1 L/min. A calibration curve using iron standards in 2% nitric acid, with varying concentrations between 12.5 ppb and 1 ppm was used to quantitate iron in protein samples.

### PFL Activation

All experiments with glycyl radical enzymes or activator enzymes were performed in a VAC Atmospheres glovebox (< 2 ppm O_2_), unless otherwise described. We activated PFL photochemically by mixing 25 μM PFL, 2.5 μM PFL-AE, 2 mM SAM, 20 mM oxamate, and 100 μM 5-deazariboflavin in activation buffer composed of 100 mM Tris, 100 mM KCl, 10 mM DTT, and 8% (w/v) glycerol at pH 7.6 in a 50 mL beaker, volumes ranging from 800 μL to 2 mL were activated each time, producing a layer of protein sample of 0.6 mm to 1.4 mm. The activation mixture was then exposed to a 1 W 405 nm LED light (Thor Labs) located approximately 5 cm above the protein sample for 1.5 h at room temperature. The lamp irradiance is 760 mW over the sample surface area. The extent of PFL activation was estimated by activity assays and EPR quantitation of the resultant G• (routinely 0.8-1.0 G•/PFL homodimer).

We measured PFL activity spectrophotometrically using a multi-enzyme assay that couples the PFL-dependent formation of acetyl-CoA from pyruvate to the production of NADH by the oxidation of malate to oxaloacetate and condensation of oxaloacetate and acetyl-CoA to citrate and CoA by malate dehydrogenase and citrate synthase, respectively, as previously reported.^56–58^ The 400 μL assay solution contained 10 mM DTT, 1 mM NAD^+^, 10 mM malate, 2 U/mL citrate synthase, 30 U/mL malate dehydrogenase and 0.05 mg/mL bovine serum albumin in 100 mM Tris buffer, adjusted to pH 8.1. Reactions were initiated by adding 3 nM aPFL (based on G•) and the rate of NADH production was calculated based on the UV absorption at 340 nm using an NADH extinction coefficient of 6.2 mM^−1^ cm^−1^ via a custom fiber-coupled Ocean Optics QEPro spectrophotometer and a DH-2000-BAL light source.^59^

### Inactivation of PFL by NO

For enzyme inactivation kinetics we buffer-exchanged aPFL into 100 mM Tris and 100 mM KCl at pH 7.6 using a Zeba Spin desalting column to a final concentration of 10 μM aPFL and maintained the solution at 30 °C using a water bath. The NONOate was then added to the aPFL solution to an estimated final concentration of either 100 μM or 1 mM from a stock solution in 100 mM glycine buffer pH 10.0. Aliquots of the reaction were sampled from 10 s to 26 min after mixing and immediately diluted 50-fold in buffer consisting of 100 mM glycine at pH 10.0 supplemented with 0.2 mg/mL of reduced deoxymyoglobin (deoxyMb) to stop the release of NO and bind any free NO. We then determined the remaining aPFL activity as described above.

To estimate the NO concentration during the decomposition of the NONOate we used deoxyMb as an NO indicator by measuring the conversion of the deoxyMb heme Soret peak shift from 431 nm to 421 nm upon binding NO.^60^ First, a solution of 10 mg/mL of reduced myoglobin was prepared by mixing oxidized myoglobin, quantitated by the heme Soret absorbance at 409 nm,^61^ with 10 mM NaDT in buffer consisting of 100 mM Tris and 100 mM KCl at pH 7.6 and then desalted to the same buffer without NaDT. The resulting deoxyMb concentration was redetermined by the heme Soret absorbance at 431 nm.^62^ To measure NO release from NONOate, 20 μM of deoxyMb was incubated with 100 μM or 1 mM NONOate in the same way as for aPFL. Aliquots were sampled during the reaction from 10 s to 26 min and immediately diluted 50-fold in buffer consisting of 100 mM glycine at pH 10.0 to stop the release of NO. NO concentration was estimated using the NO-Mb Soret absorbance of the heme-NO adduct at 421 nm.^63^

The loss of the aPFL G• upon reaction with NO was analyzed by UV-vis absorption, monitoring the characteristic G• absorption feature at 360 nm.^64^ In this assay, a solution of 55 μM aPFL in 100 mM Tris and 100 mM KCl pH 7.6 was prepared in a quartz cuvette and NONOate was added to a final concentration of 500 μM at room temperature. The reaction was followed over time by using a custom fiber-coupled Ocean Optics QEPro spectrophotometer and a DH-2000-BAL light source.

Inactivated samples were also analyzed by EPR. Similar inactivation assays were prepared using 20 μM of aPFL wt, C_418_S or C_419_S and 200 μM DEA NONOate. Samples (250 μL) were then transferred to EPR tubes and flash frozen in liquid N_2_-cooled isopentane (< −130 °C) at different reaction times and analyzed as described below.

### X-Band EPR Spectroscopy

All EPR samples were prepared in a VAC Atmosphere glovebox with < 2 ppm of O_2_ in 4 mm o.d. quartz EPR tubes and frozen in liquid N_2_-cooled isopentane (< −130 °C). EPR spectra of the samples were collected using a Bruker EMXplus EPR spectrometer at 100 K with a microwave frequency between 9.38-9.44 GHz, power of 20 μW or 2 mW, modulation amplitude of 2 G, modulation frequency of 100 kHz, time constant of 0.01 ms, scan time of 20 s, and conversion time of 16 ms. All reported spectra are the average of 30 scans. Spin quantitation was computed from the double integral of the first harmonic signal and referenced to a 4-hydroxy-TEMPO standard. All EPR spectral simulations were performed using EasySpin 6.0.0 software, and confidence intervals and standard deviations of the fitting parameters are reported.^65^

### Liquid Chromatography-Tandem Mass Spectrometry

We searched for covalent modifications of the glycine radical motif tryptic peptide in reactions of aPFL treated with NO by using trypsin digestion followed by liquid chromatography and electrospray ionization quadrupole time-of-flight tandem mass spectrometry (LC-MS/MS) on a Shimadzu LCMS-9030 q-TOF system. Samples of 25 μM aPFL were treated with 100 μM or 1 mM NONOate and incubated at room temperature for 10 min to 1 h. Reactions were stopped by mixing 5 μL of sample with 20 μL of 8 M urea in 100 mM ammonium bicarbonate and incubated for 1 h at 25 °C. Alternatively, some samples were treated with 10 mM TCEP, 10 mM DTT, or 10 mM sodium borohydride to reduce cysteines or possible nitroxyl adducts. For samples in reducing conditions cysteines were alkylated with 15 mM iodoacetamide for 60 min in the dark, and the reaction was quenched by adding 10 mM DTT or TCEP and incubated for 10 min. The samples were diluted to < 2 M urea using 100 mM ammonium bicarbonate and 0.16 μg of trypsin was added and the protein was digested at 37 °C overnight. Reactions were stopped by adding formic acid to a final concentration of 1% (v/v).

The digested samples (2-5 μg at 0.4-1 μg/μL) were injected into the LC-MS/MS. The UPLC stationary phase was a Shim-pack Arata C18 column (2.2 μm, 150 mm × 2.0 mm). The peptides were resolved by a linear gradient composed of a solution of H_2_O with 0.1% (v/v) formic acid (mobile phase A) and a solution of acetonitrile with 0.1% (v/v) formic acid (mobile phase B), from 1-30% B over 233 min, with a flow rate of 0.2 mL/min at 60 °C. Eluent from the LC was injected directly into the q-ToF. The mass spectrometer interface settings were as follow: nebulizing gas flow 2 L/min, heating gas flow 10 L/min, interface temperature 100 – 300 °C, and a desolvation temperature of 526 °C. The interface voltage was set to 4.5 V and the DL temperature to 250 °C. The spectrometer was run in data-dependent acquisition mode (DDA) with positive polarity using an event time of 0.35 s (MS^1^). Five DDA events between 200 and 1500 m/z, with a threshold of 200 counts and ions with charges between 1 and 6, were selected for fragmentation (MS^2^) using a Q1 transmission window of 1 m/z. The MS^2^ collision energy was set to 35 ± 17 V and MS^2^ ions were detected between 200 and 1,500 m/z.

The LC-MS/MS data was analyzed using the Shimadzu Lab solutions Postrun software. The MS^1^ and MS^2^ ions with different expected modifications on the glycyl radical tryptic peptide were calculated using Skyline.^66,67^ The DDA-selected MS^1^ ions were fragmented and the ions matching predicted m/z for glycyl radical tryptic peptides were manually compared to the calculated MS^2^ spectra from Skyline.

### Inactivation of PFL-AE by NO

We monitored NO inactivation of PFL-AE by measuring the effect on PFL activation (activity) and UV-vis absorbance changes associated with the essential [Fe_4_S_4_] cluster. To initiate the reaction with NO, 20 μM PFL-AE was mixed with 100 μM NONOate and incubated in a quartz cuvette for 30 min under continuous observation by UV-vis absorption. The reaction was then buffer exchanged to remove residual NO and NONOate by concentrating using 50 kDa MWCO centrifugal filters and then desalting to buffer consisting of 100 mM Tris, 100 mM KCl at pH 7.6 using a Zeba desalting column. We estimate the final concentration of NO/NONOate after these steps to be <1 μM. The resulting PFL-AE sample was then diluted 50-fold into PFL activation buffer used to activate PFL for 1 h, and PFL activity was measured using the spectrophotometric enzyme-coupled assay described above. A control sample was prepared in the same way in the absence of added NONOate.

The inactivation of PFL-AE was also monitored by EPR spectroscopy. Samples were prepared by mixing 50 μM of enzyme with 500 μM of NONOate in buffer consisting of 100 mM Tris, 100 mM KCl, 10% (w/v) glycerol adjusted to pH 7.6 and incubated at room temperature. Where indicated, samples were also prepared in the same way with the addition of 2 mM SAM or 5 mM NaDT before adding NONOate. The samples were then transferred to EPR tubes and flash frozen in liquid N_2_-cooled isopentane (< −130 °C) at different reaction times and analyzed as previously described.

### NO Treatment of Bacterial Cell Culture

To observe the effects of NO treatment in the metabolism of *E. coli*, we inoculated *E. coli* BL21 DE3 in 15 mL of LB medium and grew the cells overnight in 50 mL centrifuge tubes in a tube rotator at 20 rpm and 37 °C in anaerobic conditions in a vinyl glovebox at < 20 ppm O_2_. We used 300 μL of these cultures to inoculate 30 mL of M9 medium with 0.4% (w/v) glucose as the only carbon source and grew the cells as previously described. We took 850 μL samples over time and recorded the cell density by OD_600_. Each sample was then centrifuged at 20,000 × g for 5 min and the supernatant was used to determine the extracellular concentration of the fermentation products lactate, formate, ethanol, and acetate using spectrophotometric enzyme-coupled assays (**Supporting Information Supplemental Methods**).

The bacteriostatic and bactericidal effects of NO on anaerobically growing were assessed during growth in minimal media as previously described. At an OD_600_ of 0.35, prior to NO treatment, an aliquot of the cell culture was sampled for cell viability, reported as colony forming units (CFUs), by rapid removal from the anaerobic chamber, centrifugation and serial dilution into sterile Milli-Q water (>17 MΩ) before plating the cell culture on LB-agar plates. After collection of the pre-treatment control the cells were treated with 100 μM NONOate at OD_600_ of 0.4 and sampled again at 30 min and 4 h after NO treatment. The plated cells were then incubated at 37 ºC overnight, and colonies were counted the following day, and cell viability was determined as CFU/mL. Similar experiments were performed for cultures that were not treated with NONOate.

To analyze the effect of NO over radical species *in vivo*, and in order to observe the effect of NO on PFL and PFL-AE we transformed *E. coli* BL21 DE3 with the plasmids pCm8 wt PFL or pCal-n-EK *pfl*A and cultivated the cells overnight in 15 mL of LB medium supplemented with 50 μg/mL of chloramphenicol or kanamycin respectably. We grew the cultures overnight in 50 mL centrifuge tubes in a tube rotator at 20 rpm and 37 °C in anaerobic conditions in a vinyl glovebox at < 20 ppm O_2._ We used 300 μL of these cultures to inoculate 30 mL cultures of LB supplemented with the corresponding antibiotic. The cells were cultured to OD_600_ of 0.6 and protein expression was induced by adding 1 mM of IPTG. Induced cells were kept growing at 25 °C for 18 h, after which 100 μM or 1 mM NONOate was added and incubated for 5 min. We proceeded to collect the cells by centrifugation at 5,000 × g for 5 min and packed the cell pellet in an EPR tube and flash froze them in liquid N_2_-cooled isopentane.

### RNR Activation

We activated the class III RNR photo-chemically by mixing either 50 or 100 µM RNR, 5 or 10 µM RNR-AE, 2 mM SAM, 100 µM 5-deazariboflavin in activation buffer composed of 100 mM Tris, 100 mM KCl, 10 mM DTT and 8% (v/v) glycerol at pH 7.6 in a 50 mL beaker, for kinetic characterization or EPR analysis and NO inhibition, respectively. The mixture was exposed to the aforementioned 1 W 405 nm LED light set up for 1 h at room temperature. The extent of activation was estimated by activity assays and EPR quantitation of the resultant G• relative to a TEMPO standard. Typical activations yielded 0.01-0.04 G• per dimer.

### RNR Activity Determined via LC MS Analysis

We measured RNR activity via LC-MS analysis measuring conversion of cytidine triphosphate (CTP) to 2′-deoxycytidine triphosphate (dCTP). All activity measurements were performed in a VAC Atmosphere glove box (< 2 ppm O_2_). Following activation of 100 μM RNR, the activated RNR (aRNR) was added to a final concentration of 2 μM aRNR (50 μM total RNR) in assay buffer containing 3 mM CTP, 1 mM ATP, 10 mM formate, 30 mM KCl, 10 mM MgSO_4_, and 30 mM Tris at pH 7.6. Aliquots were taken from 15 to 300 s and quenched by boiling in a pre-heated 1.5 mL centrifuge tube in a heat block for 2 min. Quenched samples were then brought out of the anaerobic chamber and clarified via centrifugation for 15 min at 25,000 × g. 10 μL of the resulting supernatant was diluted 2-fold into dephosphorylation buffer with 1 U/mL calf alkaline phosphatase and digested for 2 h at 37 ºC according to the manufacturer’s instructions. Digestion was quenched by addition of 0.1% (v/v) TFA then diluted 20-fold with Milli-Q water and centrifuged for 30 min at 25,000 × g prior to injection on the LC-MS. Nucleo-side standards of 2′-deoxycytosine (dC, 2.5–50 μM) were prepared in Milli-Q water and adenosine (A, 12.5 μM) was used as an internal standard.

To determine the effect of NO on aRNR activity we activated 100 μM RNR and treated half of the assay (3.5 μM aRNR) with 500 μM NONOate, from a 13.6 mM stock solution in 100 mM glycine buffer pH 10.0, and incubated at room temperature for 10 min. After treatment with NO the samples were immediately diluted 5-fold in buffer consisting of 100 mM glycine at pH 10.0 to stop the release of NO. The other half of the activation assay was reserved as a control for untreated activity. The quenched NONOate-treated RNR was then diluted 2-fold into the assay buffer (10 μM RNR) and activity was measured between 0 s and 5 min as described above. The non-treated control was diluted 5-fold into assay buffer and similarly analyzed for activity from 0 s to 5 min.

Assay samples were injected onto a Shimadzu LCMS-9030 equipped with a Luna polar C18 column (1.6 μm, 50 mm × 2.1 mm). Nucleosides were resolved with a linear gradient composed of mobile phases 0.1% (v/v) TFA in water (C) and 0.1% TFA in acetonitrile (D) from 0-5% D in C over 8 min with a flow rate of 0.4 mL/min and a column temperature of 40 °C. Single ion monitoring (SIM) was used to quantitate dC (228.1 m/z ± 50 ppm) and A (268.1 m/z ± 50 ppm). Time-dependent dC formation was analyzed using the Shimadzu Lab Solutions Postrun software and determined from the chromatogram SIM MS^1^ integrated peak area normalized to the A internal standard and compared to a dC standard calibration curve.

### EPR of RNR-AE and NikJ Reacted with NO

RNR-AE and NikJ samples for EPR spectroscopy were prepared by mixing 50 μM of enzyme with 500 μM of NONOate in buffer consisting of 100 mM KCl, 100 mM Tris, and 10% (w/v) glycerol adjusted to pH 7.6 and incubated at room temperature. To examine the role of SAM and reductant on the reactions with NO, samples were prepared in the same way, but with 2 mM SAM, or by preincubating the enzyme for 15 min with 5 mM NaDT before adding NONOate. Samples (250 μL) were then transferred to EPR tubes and flash frozen in liquid N_2_-cooled isopentane at different reaction times and analyzed as described above.

## RESULTS

### Pyruvate Formate Lyase Inactivation by NO

To characterize the inactivation of aPFL by NO we used the diethylamine diazeniumdiolate (NONOate) as an NO delivery agent, which is a stable aqueous solute at pH > 9, but decomposes in a pH- and temperature-dependent first order process.^68,69^ The release of NO was quantified by monitoring the effectively diffusion-controlled and irreversible conversion of reduced myoglobin (deoxyMb) to the NO-bound myoglobin (NO-Mb) via the Soret band shift from 431 nm to 420 nm (**Supporting Information Figure S1**).^70,71^ At 30 °C and pH 7.6 we estimated a *t*_1/2_ of approximately 25 min, corresponding to an NO production rate of 2.05 μM/min and 81 μM/min at 100 μM or 1 mM NONOate, respectfully (**Figure 1**, inset). When 10 μM of aPFL is mixed with excess NONOate at either 100 μM or 1 mM and aliquots were sampled for PFL activity, activity was lost concomitantly with NO release (**Figure 1**). The activity of aPFL followed an exponential decay with an apparent rate of 0.26 μM/min at 100 μM and at 2.96 μM/min at 1 mM NONOate.

**Figure 1.**
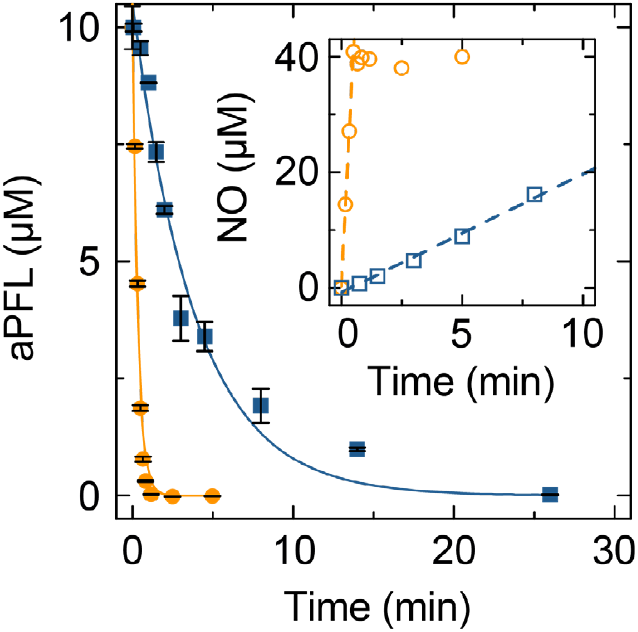
The inhibition of aPFL by NO. Samples of 10 μM aPFL were mixed with either 100 μM (blue squares) or 1 mM (orange circles) NONOate. Aliquots were taken at the indicated time points and assayed for activity. The activity decay was fit to a single exponential decay model (solid lines). Error bars represent the span of two technical replicates. Inset shows the measured NO released using deoxyMb as a NO reporter and fitted to a linear model (dashed lines).

In the measurement of residual activity of PFL following reaction with NO, enzyme and NONOate were diluted 3,000-fold and free NO was removed by binding to excess deoxyMb. Despite the elimination of NO, the enzyme remained inhibited and showed no signs of recovering over the course of the NO-free activity assay, suggesting the inhibition mechanism is irreversible. For PFL, as with all GREs, elimination of the essential G• abolishes activity, but the enzyme can be re-activated by PFL-AE, assuming no other modifications have been made. To examine the nature of NO inhibition of PFL we completely inactivated aPFL with NO, buffer exchanged the inhibited PFL into NO-free reconstitution buffer, and attempted to reactivate the enzyme via the same protocol used to generate the active enzyme initially with PFL-AE. After NO inhibition, the maximal activity that could be recovered varied between 10-20% (**Supporting Information Figure S2**). We observe a similar degree of reactivation after irreversible inactivation of aPFL by exposure to O_2_ (**Supporting Information Figure S2**), which cleaves the polypeptide chain at the glycine Cα, rendering the protein unactivatable.^37,38,72^ PFL is proposed to exhibit “half-of-sites” activation, with only one G• per homodimer, but diradical dimers have not been ruled out.^19^ As such, we attribute the observed inefficient reactivation after treatment with NO to activation of a previously non-activated monomer of a former half-of-sites dimer, or to the activation of dimers that had not been activated at all previously.

### NO Targets the Glycyl Radical of PFL

As a free radical, we expected NO to react with one of the essential amino acid radicals associated with PFL activity, namely the stable G_734_• (*E. coli* numbering) or either of the transient thiyl radicals of C_419_ or C_418_. The G• exhibits a characteristic UV-vis absorption feature at 365 nm that completely decays following exposure to NO (**Supporting Information Figure S3**). This observation is supported by X-band EPR spectroscopy, where samples were collected before addition of the NONOate and 10 min after (**Figure 2**). The characteristic asymmetric doublet EPR feature of G_734_• is completely lost after 10 min of incubation, consistent with the loss of activity.^22,25^ The UV-vis and EPR data are consistent with a mechanism of inhibition that quenches the endogenous glycyl radical on PFL. Thiyl radicals are known to react with NO to form S-nitrosothiols,^73,74^ but alkyl radicals also react with NO, thus the specific radical target of NO inhibition was not obvious.^75^ We generated C_419_S and C_418_S PFL mutants which can be reconstituted to generate G_734_•, but are completely inactive, due to the redox-inert nature of serine. While these mutants are not active, they provide insight into the reactivity of NO with PFL. Both C_419_S and C_418_S mutants showed complete radical loss by EPR over the same time frame as the wt enzyme (**Figure 2**).

**Figure 2.**
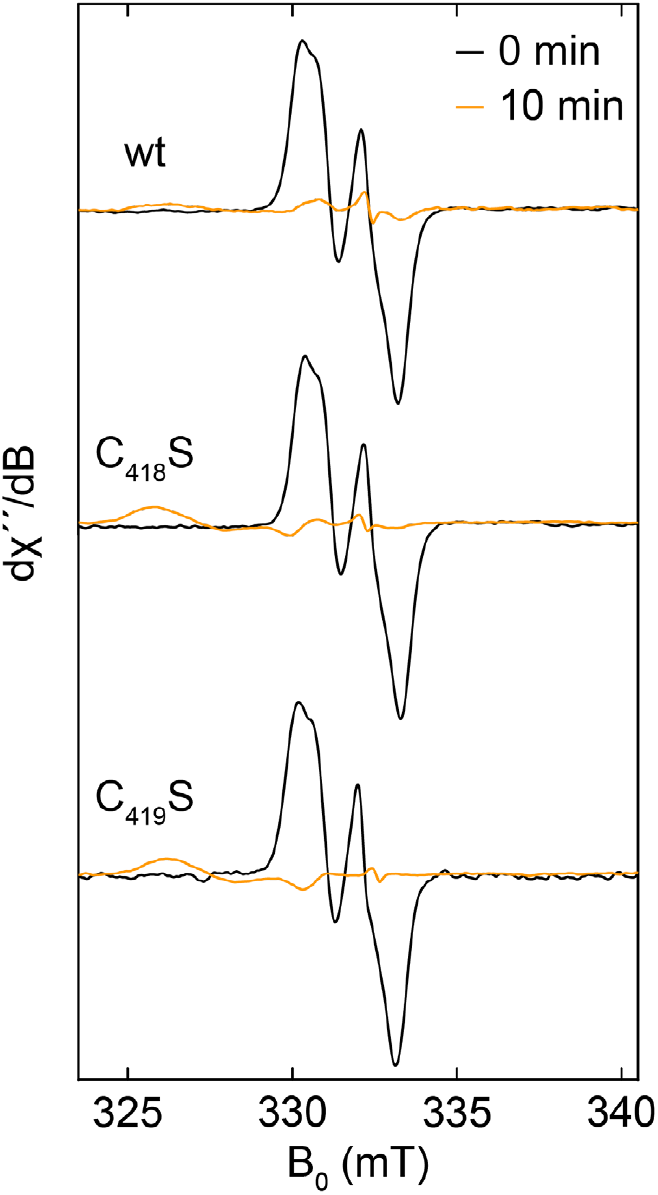
Normalized X-band EPR spectra of aPFL wt and mutants C_418_S and C_419_S before (black) and 10 mins after (orange) addition of NONOate. EPR conditions: microwave frequency, 9.3 GHz; modulation amplitude, 2 G; temperature, 100 K. 30 scans were averaged for each spectrum.

To investigate the chemical nature of the product of NO inhibition of aPFL we analyzed the product(s) of aPFL with NO by SDS-PAGE. Samples of aPFL exposed to O_2_ are cleaved at the G_734_ position, shifting the apparent electrophoretic mobility 3 kDa lower, however samples of aPFL treated with NO do not show changes in the total protein mass (**Supporting Information Figure S4**). The lack of change in apparent molecular weight by SDS-PAGE suggests that the chemical modification of aPFL does not involve cleavage of the polypeptide chain at G_734_. We also analyzed tryptic digests of the NO-inhibited PFL by LC-MS/MS for evidence of covalent modifications on peptides containing G_734_, C_419_, and C_418_. Despite an extensive analysis and the employment of reductants such as DTT, TCEP, and NaBH_4_, we observed no evidence of C_418_ or C_419_ nitrosothiols or G_734_ nitrosoalkyl/oxime or reduced amine products of any of the radical transfer peptides and no major difference in the MS^1^ chromatograms between aPFL samples and aPFL samples treated with NO, that could indicate the formation of cross-links between the modified amino acids and proximal amino acids (**Supporting Information Figure S5**). Furthermore, we observed no evidence of a C-terminal oxalylated peptide corresponding to a cleavage of the polypeptide backbone, as seen in the reaction of aPFL with O_2_.^22^ In all the samples we produced, only the unmodified NH_3_^+^-V_732_S**G**YAVR_738_-CO_2_H tryptic peptide was observed.

### NO inhibits PFL-AE

The EPR spectrum of the NO-inhibited PFL was not completely devoid of EPR-active signals (**Figure 2**). We suspected the observed residual signal originated from the reaction of NO with PFL-AE, present at 10-fold lower concentration relative to PFL in the activation reaction and carried over into the NO inhibition assay. To investigate this, we analyzed the EPR spectrum of PFL-AE in isolation before and after reacting with NO (**Figure 3**). The untreated [Fe_4_S_4_]^2+^ cluster of PFL-AE is EPR silent, whereas addition of NONOate revealed the formation of an axial EPR signal with prominent features at *g*_┴_ = 2.036 and *g*_∥_ = 2.014 (**Supporting Information Figure S6A** and **Table S2**). This EPR signature is consistent with previously characterized protein-bound monomeric dinitrosyl iron complexes (DNIC, {Fe(NO)_2_}^9^ in Enemark–Feltham notation) derived from iron-sulfur proteins.^76–80^ Spin quantification allowed to estimate a rate of DNIC formation of 8.7 μM/min, the reaction is completed in approximately 5 min with an stoichiometry of 0.8 spins/mol of PFL-AE (**Supporting Information Figure S6B** and **Table S2**). The DNIC signal did not change when PFL-AE was provided SAM, suggesting SAM binding does not protect the [Fe_4_S_4_] cluster from decomposition (**Supporting Information Figure S7** and **Table S2**). The reaction of PFL-AE and NO was also accompanied by changes in the UV-vis spectrum (**Supporting Information Figure S8A**). We observed a shift in the broad signal associated with the [Fe_4_S_4_] cluster with a peak at 410 nm to a new broad signal with a peak at 380 nm, also consistent with the monomeric DNIC.^81^ No further reaction was observed after 15 min (**Supporting Information Figure S8B**). When in the presence of the reductant NaDT, a different axial paramagnetic species was formed, with *g*_┴_ = 2.009 and *g*_∥_ = 1.970, characteristic of reduced Roussin’s red esters (**Supporting Information Figure S9A** and **Table S2**).^78,79^ As for the DNIC formation, the Roussin’s red ester signal was not affected by the presence of the substrate SAM (**Supporting Information Figure S9B** and **Table S2**). In either the presence or absence of SAM, the reduction was not complete and roughly 30% of the signal corresponds to a small signal at *g* = 2.036 remained that we attributed to non-reduced DNIC (**Supporting Information Table S2**).

**Figure 3.**
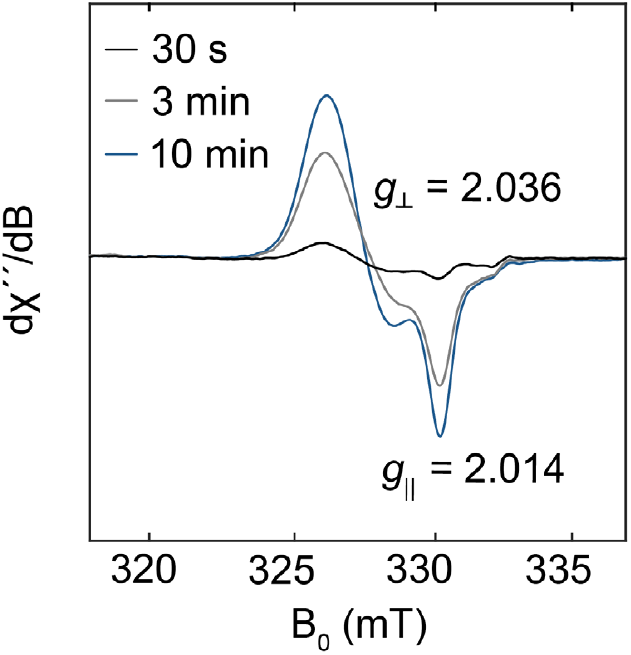
X-band EPR spectra of 50 μM PFL-AE treated with 500 μM NONOate freeze-quenched after 30 s (black), 3 mins (gray), and 10 mins (blue) of incubation at room temperature. The 10 min spectrum is plotted vs. magnetic field (B_0_, lower axis) and *g*-values are indicated. For comparison, the 30 s, 3 min and 10 min spectra are reported vs. *g* (upper axis, unlabeled). EPR conditions: microwave frequency, 9.3 GHz; modulation amplitude, 2 G; power, 2 mW; temperature, 100 K. 30 scans were averaged for each spectrum.

The [Fe_4_S_4_] cluster of PFL-AE is essential for activity, and thus, reaction with NO to form DNICs is expected to inactivate the enzyme. Indeed, reacting PFL-AE with NONOate for 30 min inhibits PFL-AE activation of PFL by 99% (**Supporting Information Figure S10**). Collectively, the results support a role of NO in degrading the essential [Fe_4_S_4_] cluster of PFL-AE via a protein-bound DNIC intermediate that renders PFL-AE inactive.

### Metabolic Consequences of Anaerobic *E. coli* NO Exposure

Having established the *in vitro* irreversible inactivation of PFL and PFL-AE, we sought to investigate the *in vivo* consequences of NO treatment on the metabolism of *E. coli*. We cultivated *E. coli* cells anaerobically in minimal media with glucose as the unique carbon source, directing glucose metabolism to anaerobic fermentation. Cells in mid-exponential phase were treated with 100 μM NONOate or left unperturbed as a control (**Figure 4** and **Supporting Information Figure S11**). The introduction of NO immediately inhibited cell growth for a period of more than 7 h, demonstrating a bacteriostatic effect of NO in anaerobic conditions. NO also displayed bactericidal effects; cells in the mid-exponential phase displayed a significant decrease in cell viability by 65% within 30 min and 77% after 4 h of NO treatment (**Supporting Information Figure S12**). We also measured the extracellular concentration of the fermentation products lactate, formate, acetate, and ethanol using enzyme-coupled assays (**Supporting Information Figure S13**). Non-treated cultures produce principally formate and lower concentrations of ethanol and acetate, however the addition of NO causes a complete inhibition of the production of ethanol, formate, and acetate and the cell culture accumulated lactate at higher rates and concentrations than in the non-treated cultures. This metabolic shift suggests the *in vivo* inhibition of PFL and PFL-AE by NO.

**Figure 4.**
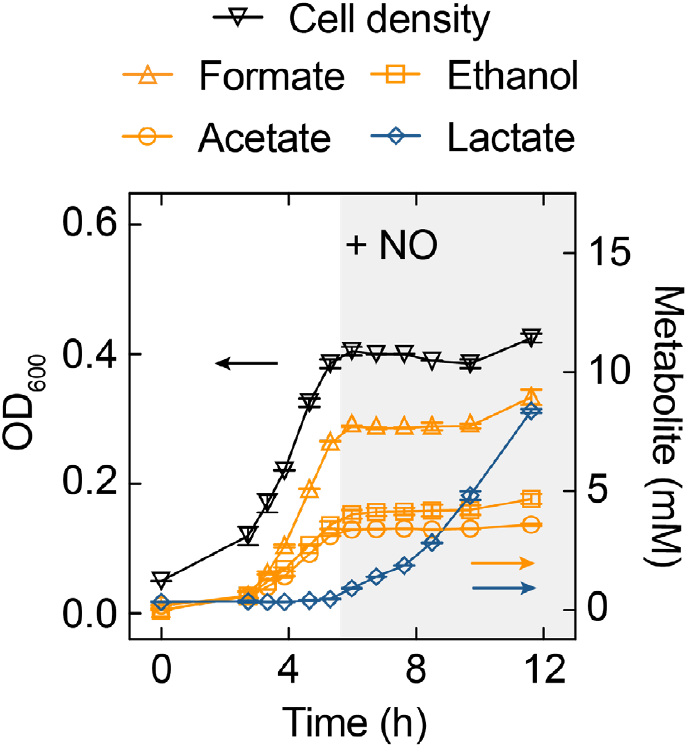
The metabolic impact of NO on anaerobically growing *E. coli*. Anaerobic cultures of *E. coli* were subjected to treatment with 100 μM NONOate at mid-exponential phase (gray shaded area). Cell density was estimated from the OD_600_ (black triangles) and extracellular concentration of the fermentation products lactate (blue diamonds), acetate (orange circles), ethanol (orange squares), and formate (orange triangles) were determined using enzymatic coupled assays. Error bars represent the span between two biological replicates.

To compare the mechanism of PFL and PFL-AE inhibition *in vitro* with the potentially more complex chemistry *in vivo*, we attempted whole-cell EPR of anaerobically grown *E. coli* overexpressing either PFL or PFL-AE (**Figure 5**). PFL and PFL-AE can be overexpressed in anaerobic conditions and represent >10% of the total cell protein content, as observed by SDS-PAGE (**Supporting Information Figure S14**). After protein induction and subsequent treatment with NO, we harvested the cells and analyzed them by EPR. *E. coli* over-expressing PFL can reconstitute the PFL glycyl radical, which can readily be observe by X-band EPR (**Figure 5A**, inset). The addition of NO quenches the PFL glycyl radical on a timescale similar to the one observed *in vitro*. We also observed significant DNIC formation in these samples, with characteristic features at *g*_┴_ = 2.037 and *g*_∥_ = 2.015, despite no over-expression of PFL-AE (**Figure 5A, Supporting Information Figure S15**, and **Table S3**).

**Figure 5.**
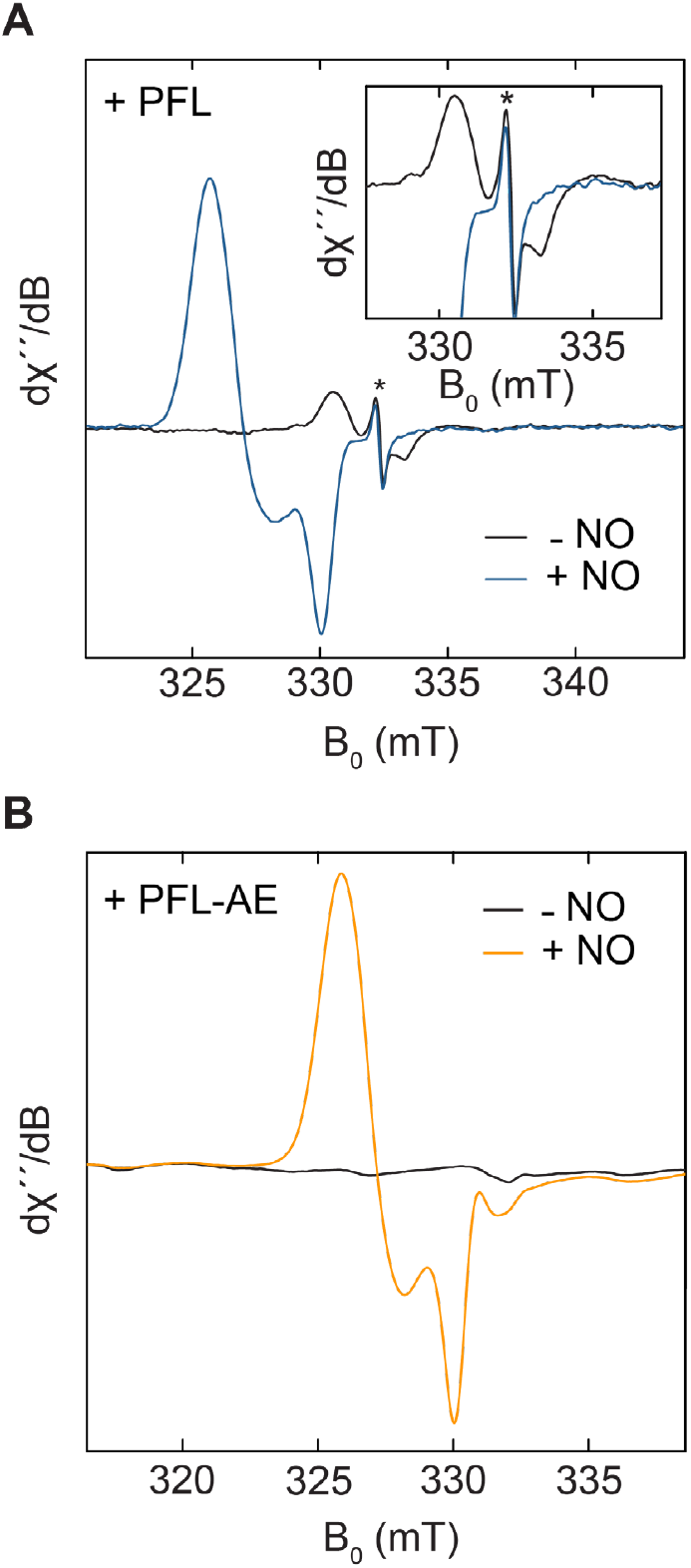
X-band EPR spectra of anaerobic *E. coli* cultures over-expressing **A** PFL or **B** PFL-AE before (black) and after (blue and orange, respectively) exposure to 100 μM NONOate for 10 min. Inset in A shows an expanded view of the G• signal. The asterisk “*” indicates a cavity artifact. EPR conditions: microwave frequency, 9.3 GHz; modulation amplitude, 2 G; power, 20 μW (A) or 2 mW (B), temperature, 100 K. 30 scans were averaged for each spectrum.

In *E. coli* cultures over-expressing PFL-AE, a glycyl radical signal can also be observed at lower intensity than the signal observed in samples of the PFL over-expressing cultures (**Supporting Information Figure S16A**). This signal can be attributed to the genomic PFL expression that can be as high as 20 μM in the cytoplasm of anaerobically growing *E. coli*.^20,23^ However, this glycyl radical signal could correspond to other glycyl radical enzymes in the *E. coli* proteome, such as the class III RNR, ketobutyrate formate lyase, YbiW, or PflD.^24,82^ The [Fe_4_S_4_]^2+^ of PFL-AE is EPR silent before treatment with NO *in vivo*. Following treatment with 100 μM NONOate, a signal consistent with the formation of DNICs appears with the disappearance of the low-intensity glycyl radical signal (**Figure 5B** and **Supporting Information Figure S16**), and treatment of the cultures with 1 mM of NONOate produced a composite signal of DNIC and Roussin’s red ester (**Supporting Information Figure S16B and Supporting Information Table S3**). Non-transformed *E. coli* cells exposed to NO produce the same signal, although at lower intensity but with similar *g*-values (**Supporting Information Figure S17** and **Table S3**). We attribute this signal to the formation of DNICs in other iron-sulfur proteins present in the *E. coli* proteome.

### NO Inhibits a Broad Range of GREs and Their Activases

Our observation of the NO effect on *E. coli* metabolism and loss of the whole-cell glycyl radical EPR signal suggests the *in vivo* inhibition of PFL. Several enzymes in the glycyl radical enzyme family are associated with primary metabolism in *E. coli*. Therefore, we sought to investigate the reactivity of NO with the *E. coli* class III RNR and its radical SAM activator enzyme, RNR-AE. The class III RNR is a glycyl radical enzyme found in obligate and facultative anerobic bacteria and archaea, including *E. coli*, and is essential for the anaerobic reduction of nucleotides to deoxynucleotides for *de novo* DNA synthesis and repair.^83,84^

As with PFL, we examined whether the glycyl radical of RNR reacts with exogenous NO, inhibiting the enzyme. We exposed activated RNR (aRNR) to NO, analogous to our prior experiments with aPFL, and followed the reaction by EPR. After 10 min of treatment with 500 µM NONOate we observed a complete loss of the aRNR glycyl radical signature, as well as the formation of a DNIC signal we attribute to the reaction of remnant RNR-AE with NO from the RNR activation mixture (**Figure 6**). To investigate the effect of glycyl radical loss on the activity of aRNR we utilized LC-MS to quantitate the dephosphorylated reaction product, dC, of CTP reduction (**Supporting Information Figure S18** and **S19**). Non-treated aRNR produced dCTP with a linear rate over 1 min which ultimately slowed as the reaction approached completion; displaying a specific activity of 1,500 ± 100 nmol/min×mg, consistent with literature values (**Supporting Information Figure S19**).^50^ On the other hand, aRNR treated with NONOate for 10 min showed no detectable dC formation over 5 min in the activity assay (<3.5 nmol/min×mg, **Figure 6** inset and **Supporting Information Figure S19**).

**Figure 6.**
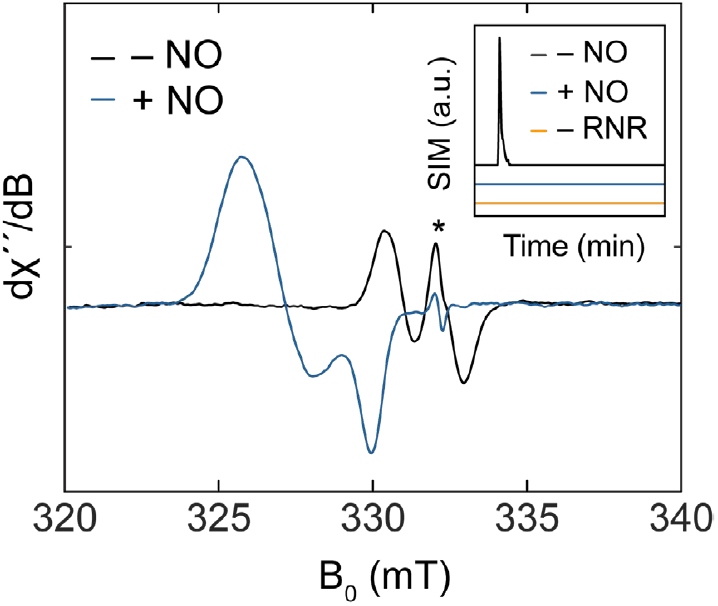
X-band EPR spectra of 100 μM reconstituted *E. coli* class III RNR before (black) and after (blue) treatment with 500 μM NONOate for 10 min. The asterisk “*” indicates a cavity artifact. EPR conditions: microwave frequency, 9.3 GHz; modulation amplitude, 2 G; power, 20 μW; temperature, 100 K. 30 scans were averaged for each spectrum. Inset shows dC production by aRNR over 5 min. Single ion monitoring (SIM) LC-MS for dC was performed for aRNR (black), aRNR treated with 500 μM NONOate for 10 min (blue), or assay buffer without aRNR (orange).

To examine the reaction of NO with the RNR-AE in greater detail, we also reacted RNR-AE with NO in isolation and followed the reaction by EPR. As expected, upon reaction with NO, RNR-AE produced a similar DNIC EPR signal to that observed for PFL-AE (**Supporting Information Figure S20A**), with similar *g*_┴_ = 2.035 and *g*_∥_ = 2.014 (**Supporting Information Figure S20B** and **Table S4**) to PFL-AE, and an estimated rate of formation of 16.3 μM/min, again similar to the one observed for PFL-AE (**Supporting Information Figure S20C**). As for PFL-AE, the DNIC signal of RNR-AE and its *g*-values were not affected by the presence of the substrate SAM (**Supporting Information Figure S21** and **Table S4**). In the presence of NO and reducing conditions, RNR-AE formed the characteristic axial paramagnetic signal of Roussin’s red ester, with *g*_┴_ = 2.010 and *g*_∥_ = 1.971, and 35% of the signal corresponding to non-reduced DNIC (**Supporting Information Figure S22** and **Table S4**). The results of both NO inhibition of the class III RNR and the associated RNR-AE suggest a general mechanism of glycyl radical and activator enzyme inhibition by NO.

To further establish the generality of inhibition of radical SAM enzyme superfamily members by exogenous NO, we analyzed the inhibition of the enzyme NikJ, a radical SAM enzyme that catalyzes the C5′ extension from enoylpyruvyluridine monophosphate (EP-UMP) to octosyl acid in the biosynthesis pathway of nikkomycins.^85,86^ PFL-AE and RNR-AE are homologs and share similar function and sequence (sequence identity 28%), while NikJ does not share significant similarity with either PFL-AE or RNR-AE, representing a distant homolog from the same superfamily. The reaction of NikJ with NO resulted in the formation of an EPR-active DNIC species and followed slower kinetics relative to PFL-AE and RNR-AE (**Supporting Information Figure S23** and **Table S5**). As observed for PFL-AE and RNR-AE the presence of the substrate SAM does not change the *g*-values of the observed signal (**Supporting Information Figure S24** and **Table S5**). NikJ reduced with NaDT produced EPR signals consistent with both a DNIC and Roussin’s red ester (**Supporting Information Figure S25A** and **Table S5**), while in the presence of SAM, a signal consistent with Roussin’s red ester was observed with nearly identical spectral features as those of PFL-AE and RNR-AE, and showing low concentrations of a non-reduced DNIC signal (**Supporting Information Figure S25B** and **Table S5**).

## DISCUSSION

Phagocytes, including macrophages, microglia and neutrophils, along with intestinal epithelial cells, respond to proinflammatory cytokines by expressing the inducible nitric oxide synthase (iNOS).^87–89^ iNOS catalyzes the oxidation of L-arginine into L-citrulline and NO, releasing high concentrations of NO into sites of infection. NO acts as both a bactericidal and bacteriostatic agent and its mechanisms of action have been investigated in both *in vivo*^39,90,91^ and *in vitro*^92,93^ models. Many molecular targets of NO toxicity have been recognized in bacteria, principally in aerobic conditions; NO inhibits DNA replication by targeting DNA-binding zinc metalloproteins^94^ and deoxynucleotides production by reacting with cysteinyl and tyrosyl radicals of RNR.^95,96^ Additionally, NO impairs respiration by binding to the Cu_B_ of cytochrome bo^97^ and central metabolism by inhibiting many enzymes in the glycolytic pathway and the TCA cycle.^80,98^ However, in anaerobic conditions, relevant to gastrointestinal infections, the molecular targets of NO have not been identified with molecular specificity. This knowledge gap limits our understanding of the innate immune response in anaerobic environments and hinders the potential application of NO as a therapeutic agent in treating gastrointestinal infections.^17,99^

In this study, we identified PFL as a direct target of NO and characterized the mechanism of inhibition. Upon exposure to NO, PFL is inhibited with an apparent diffusion-controlled rate, suggesting PFL is a kinetically important microbial metabolic target of NO. Thiyl radicals react with NO at diffusion-controlled rates,^74^ however our EPR and site-directed mutagenesis data following radical quenching strongly suggest that NO reacts directly with the glycyl radical of the active enzyme. In PFL, C• and G• exist in an equilibrium favoring the formation of G•.^100^ The lack of evidence of nitrosothiol formation during aPFL inhibition implies a rapid rate of reaction with G• and a low concentration of C• in the G• ⇄ C• equilibrium. Despite our inability to directly demonstrate the nature of the NO adduct(s) to PFL by LC-MS/MS, we hypothesize a radical-radical coupling mechanism. The lack of observation of an NO adduct on Gly_734_ may be related to instability of the modification during MS sample preparation or instability during ionization in the mass spectrometer; such effects have previously been observed for cysteinyl S-NO modified peptides, and their detection generally requires the use of indirect techniques.^101–103^ Together, these data demonstrate that NO is a potent inhibitor of the glycyl radical enzyme PFL via the quenching of the essential glycyl radical cofactor.

Studies on the effect of NO on anaerobic gastrointestinal bacterial communities indicate that NO exposure influences fermentation products and community composition.^104^ To evaluate the effect of PFL inactivation by NO on anaerobic metabolism in a model gastrointestinal microbe, we examined the effect of NO on anaerobic *E. coli* cell growth, metabolism, and enzyme post-translational modifications. Using whole-cell EPR, we show that NO reacts with glycyl radicals and FeS clusters in *E. coli* cultures at a concentration of 100 μM−comparable to NO levels measured in gastrointestinal samples from patients with inflammatory conditions such as ulcerative colitis and Crohn’s disease.^10,12^ Additionally, we observe that NO causes metabolic changes in anaerobic *E. coli* cultures, arresting the production of the metabolites formate, acetate, and ethanol and halting cell growth and viability. In the absence of PFL activity, lactate accumulates as the product of pyruvate metabolism via lactate dehydrogenase, which is not inhibited by NO.^39^ Similar effects have been reported in anaerobic fermentation from healthy human fecal samples treated with NO, where NO produced a long-lasting impact on the metabolome.^104^ In *E. coli*, deletion of the *pflB* gene shifts metabolism to lactate formation, similar to *E. coli* treated with NO, but does not completely impair growth.^105,106^ Our observation that growth is effectively halted despite continued lactate production implies a substantial maintenance and repair burden associated with NO exposure. We hypothesized that the inhibition of class III RNR, another glycyl radical enzyme essential to *de novo* deoxyribonucleotide synthesis, may contribute to the bacteriostatic effect of NO while maintaining lactate fermentation. The glycyl radical enzyme, RNR, was indeed inhibited by NO *in vitro*, with a reaction rate similar to PFL, suggesting DNA replication and repair are also inhibited, yet further experiments are required to determine the consequences of RNR inhibition by NO *in vivo*.

We further demonstrate that NO inactivates PFL-AE and RNR-AE. Therefore, NO acts as a general GRE-AE inhibitor, serving a two-fold mechanism of inactivating GRE chemistry, at both the level of GREs and GRE-AEs. Our EPR data evidences the NO reactivity with the enzymes [Fe_4_S_4_] cluster and the formation of DNICs, with similar spectroscopic properties to previously reported iron-sulfur cluster proteins and enzymes.^76–80^ The DNIC formation is accompanied by a complete loss of activity for PFL-AE, as the [Fe_4_S_4_] is essential for enzyme activity. Radical SAM enzyme iron-sulfur clusters are coordinated by three cysteines and, when present, the amino and carboxy group of SAM.^27^ The presence of SAM provided no protection from NO inhibition for either PFL-AE or RNR-AE, nor the distantly related radical SAM enzyme NikJ, demonstrating that SAM binding and coordination of the fourth iron does not protect the iron-sulfur cluster from decomposition.

The collective evidence that GREs and radical SAM enzymes, including GRE-AEs, are irreversibly inhibited by NO. This suggests that the mechanism by which NO inhibits GREs and GRE-AEs is part of the innate immune response, triggered by gastrointestinal pathogens. Many metabolic pathways in the gut microbiome involve GREs or radical SAM enzymes, indicating that NO effects may be extensive.^21,24,27,107–111^ Additionally, the enzyme pyruvate:ferredoxin oxidoreductase is also involved in anaerobic metabolism of pyruvate and harbors an iron-sulfur cluster, and is another likely target of anaerobic metabolic inhibition by NO.^112^ What remains to be seen is whether the NO response differentially affects the microbiome community, and what defense mechanisms may have adapted to anaerobic NO exposure that may inform therapeutic approaches aimed at enhancing the innate immune response.

## ASSOCIATED CONTENT

### Supporting Information

Supplemental methods, list of primers, UV/vis spectroscopy of myoglobin as an NO sensor, PFL reactivation, aPFL NO inactivation followed by UV/vis spectroscopy, Inactivation of aPFL followed by SDS-PAGE, peptide LC-MS/MS of aPFL inhibited with NO, EPR characterization of PFL-AE and kinetics, EPR simulations parameters for PFL-AE treated with NO, EPR characterization of PFL-AE reacted with NO in the presence of SAM, PFL-AE NO inactivation followed by UV/Vis spectroscopy, EPR characterization of reduced PFL-AE with NO, Inhibition of PFL-AE activity by NO, metabolic analysis of anaerobically growing *E. coli*, metabolites calibration curves, over-expression of PFL and PFL-AE in anaerobic conditions, EPR characterization of whole cells overexpressing PFL or PFL-AE treated with NO, EPR characterization of RNR-AE NO inactivation, EPR simulations parameters for RNR-AE treated with NO, EPR characterization of reduced RNR-AE reacted with NO, deoxycytidine calibration curve, LC-MS activity assay of aRNR and NO treated aRNR, EPR characterization of NikJ NO inactivation, EPR characterization of reduced NikJ with NO and EPR simulations parameters for NikJ treated with NO.

## AUTHOR INFORMATION

**Author Contributions**

The manuscript was written through contributions of all authors. All authors have given approval to the final version of the manuscript.

## ACKNOWLEDGMENTS

We gratefully acknowledge Prof. Joan Broderick for providing the pCAL-n-EK plasmid. We also acknowledge Dr. Squire Booker for providing the plasmid pDB1282 and helpful discussion. We also thank Dr. Peter Ford for stimulating conversations regarding NO delivery. The plasmid TEVSH was a gift from Helena Berglund (AddGene plasmid #125194). The plasmids pFGET19_Ulp1 and pHYRSF53 were a gift from Hideo Iwai (Addgene plasmid #64697 and #64696). The research reported here used shared facilities of the UC Santa Barbara Materials Research Science and Engineering Center (MRSEC, NSF DMR-1720256), a member of the Materials Research Facilities Network (http://www.mrfn.org). The Department of Chemistry and Biochemistry Mass Spectrometry Facility instrumentation was supported by the Department of Defense DURIP grant number N00014-23-1-2197. J.C.C. thanks Anid/Subdirección De Capital Humano/Beca de Doctorado Becas Chile/72200442 for their support. B.L.G thanks the National Institute of General Medical Sciences (R35GM154993), the Hellman Faculty Fellowship, and a UCSB Faculty Research Grant for funding. Additionally, the authors acknowledge the contributions of undergraduate researchers Lindsey Calva, Aisling C. Parast, and Maclean Thomson.

---

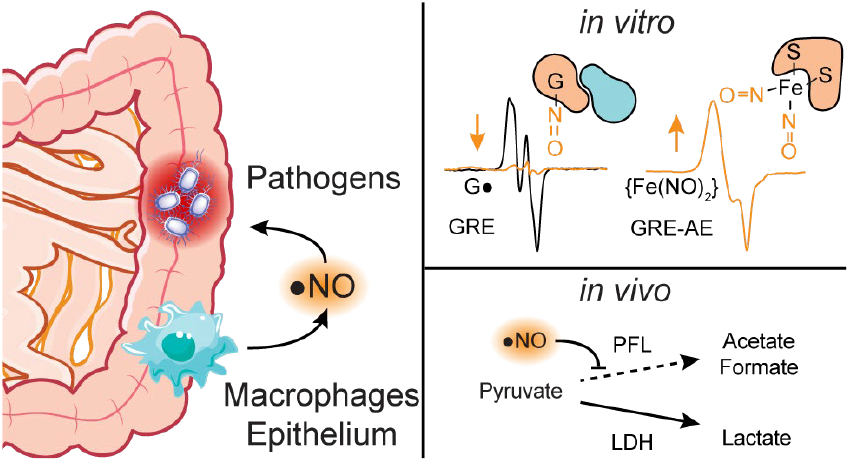

## REFERENCES

(1) Kotloff, K. L.; Riddle, M. S.; Platts-Mills, J. A.; Pavlinac, P.; Zaidi, A. K. M. Shigellosis. Lancet 2018, 391 (10122), 801–812. DOI: 10.1016/S0140-6736(17)33296-8.

(2) Yamamoto, N.; Droffner, M. L. Mechanisms Determining Aerobic or Anaerobic Growth in the Facultative Anaerobe Salmonella Typhimurium. Proc. Natl. Acad. Sci. U.S.A. 1985, 82 (7), 2077–2081. DOI: 10.1073/pnas.82.7.2077.

(3) Nataro, J. P.; Kaper, J. B. Diarrheagenic Escherichia Coli. Clin. Microbiol. Rev. 1998, 11 (1), 142–201. DOI: 10.1128/cmr.11.1.142.

(4) Schobert, M.; Jahn, D. Anaerobic Physiology of Pseudomonas Aeruginosa in the Cystic Fibrosis Lung. Int. J. Med. Microbiol. 2010, 300 (8), 549–556. DOI: 10.1016/j.ijmm.2010.08.007.

(5) Hall, J. W.; Ji, Y. Sensing and Adapting to Anaerobic Conditions by Staphylococcus Aureus. In Advances in Applied Microbiology; Sariaslani, S., Gadd, G. M., Eds.; Academic Press, 2013; Vol. 84, pp 1–25. DOI: 10.1016/B978-0-12-407673-0.00001-1.

(6) Million, M.; Diallo, A.; Raoult, D. Gut Microbiota and Malnutrition. Microb. Pathog. 2017, 106, 127–138. DOI: 10.1016/j.micpath.2016.02.003.

(7) Million, M.; Raoult, D. Linking Gut Redox to Human Microbiome. Hum. Microbiome J. 2018, 10, 27–32. DOI: 10.1016/j.humic.2018.07.002.

(8) Pircalabioru, G.; Aviello, G.; Kubica, M.; Zhdanov, A.; Paclet, M.-H.; Brennan, L.; Hertzberger, R.; Papkovsky, D.; Bourke, B.; Knaus, U. G. Defensive Mutualism Rescues NADPH Oxidase Inactivation in Gut Infection. Cell Host Microbe 2016, 19 (5), 651–663. DOI: 10.1016/j.chom.2016.04.007.

(9) Fang, F. C. Antimicrobial Reactive Oxygen and Nitrogen Species: Concepts and Controversies. Nat. Rev. Microbiol. 2004, 2 (10), 820–832. DOI: 10.1038/nrmicro1004.

(10) Lundberg, J. O. N.; Lundberg, J. M.; Alving, K.; Hellström, P. M. Greatly Increased Luminal Nitric Oxide in Ulcerative Colitis. Lancet 1994, 344 (8938), 1673–1674. DOI: 10.1016/S0140-6736(94)90460-X.

(11) Kubes, P.; McCafferty, D.-M. Nitric Oxide and Intestinal Inflammation. Am. J. Med. 2000, 109 (2), 150–158. DOI: 10.1016/S0002-9343(00)00480-0.

(12) Kimura, H.; Miura, S.; Shigematsu, T.; Ohkubo, N.; Tsuzuki, Y.; Kurose, I.; Higuchi, H.; Akiba, Y.; Hokari, R.; Hirokawa, M.; Serizawa, H.; Ishii, H. Increased Nitric Oxide Production and Inducible Nitric Oxide Synthase Activity in Colonic Mucosa of Patients with Active Ulcerative Colitis and Crohn’s Disease. Dig. Dis. Sci. 1997, 42 (5), 1047–1054. DOI: 10.1023/A:1018849405922.

(13) Mu, K.; Yu, S.; Kitts, D. D. The Role of Nitric Oxide in Regulating Intestinal Redox Status and Intestinal Epithelial Cell Functionality. Int. J. Mol. Sci. 2019, 20 (7), 1755. DOI: 10.3390/ijms20071755.

(14) Katsube, T.; Tsuji, H.; Onoda, M. Nitric Oxide Attenuates Hydrogen Peroxide-Induced Barrier Disruption and Protein Tyrosine Phosphorylation in Monolayers of Intestinal Epithelial Cell. Biochim. Biophys. Acta, Mol. Cell Res. 2007, 1773 (6), 794–803. DOI: 10.1016/j.bbamcr.2007.03.002.

(15) Vázquez-Torres, A.; Fang, F. C. Nitric Oxide in Salmonella and Escherichia Coli Infections. EcoSal Plus 2005, 1 (2). DOI: 10.1128/ecosalplus.8.8.8.

(16) Porrini, C.; Ramarao, N.; Tran, S.-L. Dr. NO and Mr. Toxic – the Versatile Role of Nitric Oxide. Biol. Chem. 2020, 401 (5), 547– 572. DOI: 10.1515/hsz-2019-0368.

(17) Rouillard, K. R.; Novak, O. P.; Pistiolis, A. M.; Yang, L.; Ahonen, M. J. R.; McDonald, R. A.; Schoenfisch, M. H. Exogenous Nitric Oxide Improves Antibiotic Susceptibility in Resistant Bacteria. ACS Infect. Dis. 2021, 7 (1), 23–33. DOI: 10.1021/acsinfecdis.0c00337.

(18) Rouillard, K. R.; Hill, D. B.; Schoenfisch, M. H. Antibiofilm and Mucolytic Action of Nitric Oxide Delivered via Gas or Macromolecular Donor Using in Vitro and Ex Vivo Models. J. Cyst. Fibros. 2020, 19 (6), 1004–1010. DOI: 10.1016/j.jcf.2020.03.004.

(19) Knappe, J.; Schacht, J.; Möckel, W.; Höpner, Th.; Vetter Jr., H.; Edenharder, R. Pyruvate Formate-Lyase Reaction in Escherichia Coli. Eur. J. Biochem. 1969, 11 (2), 316–327. DOI: 10.1111/j.1432-1033.1969.tb00775.x.

(20) Knappe, J.; Sawers, G. A Radical-Chemical Route to Acetyl-CoA: The Anaerobically Induced Pyruvate Formate-Lyase System of Escherichia Coli. FEMS Microbiol. Rev. 1990, 6 (4), 383–398. DOI: 10.1111/j.1574-6968.1990.tb04108.x.

(21) Levin, B. J.; Huang, Y. Y.; Peck, S. C.; Wei, Y.; Campo, A. M.; Marks, J. A.; Franzosa, E. A.; Huttenhower, C.; Balskus, E. P. A Prominent Glycyl Radical Enzyme in Human Gut Microbiomes Metabolizes Trans-4-Hydroxy-l-Proline. Science 2017, 355 (6325). DOI: 10.1126/science.aai8386.

(22) Wagner, A. F.; Frey, M.; Neugebauer, F. A.; Schäfer, W.; Knappe, J. The Free Radical in Pyruvate Formate-Lyase Is Located on Glycine-734. Proc. Natl. Acad. Sci. U.S.A. 1992, 89 (3), 996–1000. DOI: 10.1073/pnas.89.3.996.

(23) Crain, A. V.; Broderick, J. B. Pyruvate Formate-Lyase and Its Activation by Pyruvate Formate-Lyase Activating Enzyme. J. Biol. Chem. 2014, 289 (9), 5723–5729. DOI: 10.1074/jbc.M113.496877.

(24) Backman, L. R. F.; Funk, M. A.; Dawson, C. D.; Drennan, C. L. New Tricks for the Glycyl Radical Enzyme Family. Crit. Rev. Biochem. Mol. Biol. 2017, 52 (6), 674–695. DOI: 10.1080/10409238.2017.1373741.

(25) Knappe, J.; Neugebauer, F. A.; Blaschkowski, H. P.; Gänzler, M. Post-Translational Activation Introduces a Free Radical into Pyruvate Formate-Lyase. Proc. Natl. Acad. Sci. U.S.A. 1984, 81 (5), 1332–1335. DOI: 10.1073/pnas.81.5.1332.

(26) Broderick, J. B.; Duderstadt, R. E.; Fernandez, D. C.; Wojtuszewski, K.; Henshaw, T. F.; Johnson, M. K. Pyruvate Formate-Lyase Activating Enzyme Is an Iron-Sulfur Protein. J. Am. Chem. Soc. 1997, 119 (31), 7396–7397. DOI: 10.1021/ja9711425.

(27) Broderick, J. B.; Duffus, B. R.; Duschene, K. S.; Shepard, E. M. Radical S-Adenosylmethionine Enzymes. Chem. Rev. 2014, 114 (8), 4229–4317. DOI: 10.1021/cr4004709.

(28) Sawers, R. G.; Clark, D. P. Fermentative Pyruvate and Acetyl-Coenzyme A Metabolism. EcoSal Plus 2004, 1 (1), 10.1128/ecosalplus.3.5.3. DOI: 10.1128/ecosalplus.3.5.3.

(29) Pinske, C.; Sawers, R. G. Anaerobic Formate and Hydrogen Metabolism. EcoSal Plus 2016, 7 (1), 10.1128/ecosalplus.esp-0011-2016. DOI: 10.1128/ecosalplus.esp-0011-2016.

(30) Mejillano, M. R.; Jahansouz, H.; Matsunaga, T. O.; Kenyon, G. L.; Himes, R. H. Formation and Utilization of Formyl Phosphate by N10-Formyltetrahydrofolate Synthetase: Evidence for Formyl Phosphate as an Intermediate in the Reaction. Biochemistry 1989, 28 (12), 5136–5145. DOI: 10.1021/bi00438a034.

(31) Mulliez, E.; Ollagnier, S.; Fontecave, M.; Eliasson, R.; Reichard, P. Formate Is the Hydrogen Donor for the Anaerobic Ribonucleotide Reductase from Escherichia Coli. Proc. Natl. Acad. Sci. U.S.A. 1995, 92 (19), 8759–8762. DOI: 10.1073/pnas.92.19.8759.

(32) Abernathy, J.; Corkill, C.; Hinojosa, C.; Li, X.; Zhou, H. Deletions in the Pyruvate Pathway of Salmonella Typhimurium Alter SPI1-Mediated Gene Expression and Infectivity. J. Anim. Sci. Biotechnol. 2013, 4 (1), 5. DOI: 10.1186/2049-1891-4-5.

(33) Kuntumalla, S.; Zhang, Q.; Braisted, J. C.; Fleischmann, R. D.; Peterson, S. N.; Donohue-Rolfe, A.; Tzipori, S.; Pieper, R. In Vivo versus in Vitro Protein Abundance Analysis of Shigella Dysenteriae Type 1 Reveals Changes in the Expression of Proteins Involved in Virulence, Stress and Energy Metabolism. BMC Microbiol. 2011, 11 (1), 147. DOI: 10.1186/1471-2180-11-147.

(34) Neumann-Schaal, M.; Jahn, D.; Schmidt-Hohagen, K. Metabolism the Difficile Way: The Key to the Success of the Pathogen Clostridioides Difficile. Front. Microbiol. 2019, 10. DOI: 10.3389/fmicb.2019.00219.

(35) Yesilkaya, H.; Spissu, F.; Carvalho, S. M.; Terra, V. S.; Homer, K. A.; Benisty, R.; Porat, N.; Neves, A. R.; Andrew, P. W. Pyruvate Formate Lyase Is Required for Pneumococcal Fermentative Metabolism and Virulence. Infect. Immun. 2009, 77 (12), 5418–5427. DOI: 10.1128/iai.00178-09.

(36) Merriman, J. A.; Xu, W.; Caparon, M. G. Central Carbon Flux Controls Growth/Damage Balance for Streptococcus Pyogenes. PLoS Pathog. 2023, 19 (6), e1011481. DOI: 10.1371/journal.ppat.1011481.

(37) Reddy, S. G.; Wong, K. K.; Parast, C. V.; Peisach, J.; Magliozzo, R. S.; Kozarich, J. W. Dioxygen Inactivation of Pyruvate Formate-Lyase: EPR Evidence for the Formation of Protein-Based Sulfinyl and Peroxyl Radicals. Biochemistry 1998, 37 (2), 558–563. DOI: 10.1021/bi972086n.

(38) Zhang, W.; Wong, K. K.; Magliozzo, R. S.; Kozarich, J. W. Inactivation of Pyruvate Formate-Lyase by Dioxygen: Defining the Mechanistic Interplay of Glycine 734 and Cysteine 419 by Rapid Freeze-Quench EPR. Biochemistry 2001, 40 (13), 4123–4130. DOI: 10.1021/bi002589k.

(39) Richardson, A. R.; Libby, S. J.; Fang, F. C. A Nitric Oxide–Inducible Lactate Dehydrogenase Enables Staphylococcus Aureus to Resist Innate Immunity. Science 2008, 319 (5870), 1672–1676. DOI: 10.1126/science.1155207.

(40) Palmieri, E. M.; Gonzalez-Cotto, M.; Baseler, W. A.; Davies, L. C.; Ghesquière, B.; Maio, N.; Rice, C. M.; Rouault, T. A.; Cassel, T.; Higashi, R. M.; Lane, A. N.; Fan, T. W.-M.; Wink, D. A.; McVicar, D. W. Nitric Oxide Orchestrates Metabolic Rewiring in M1 Macrophages by Targeting Aconitase 2 and Pyruvate Dehydrogenase. Nat. Commun. 2020, 11 (1), 698. DOI: 10.1038/s41467-020-14433-7.

(41) Gardner, P. R.; Costantino, G.; Salzman, A. L. Constitutive and Adaptive Detoxification of Nitric Oxide in Escherichia Coli: Role of Nitric-Oxide Dioxygenase in the Protection of Aconitase. J. Biol. Chem. 1998, 273 (41), 26528–26533. DOI: 10.1074/jbc.273.41.26528.

(42) Welter, R.; Yu, L.; Yu, C.-A. The Effects of Nitric Oxide on Electron Transport Complexes. Arch. Biochem. Biophys. 1996, 331 (1), 9–14. DOI: 10.1006/abbi.1996.0276.

(43) Broderick, J. B.; Henshaw, T. F.; Cheek, J.; Wojtuszewski, K.; Smith, S. R.; Trojan, M. R.; McGhan, R. M.; Kopf, A.; Kibbey, M.; Broderick, W. E. Pyruvate Formate-Lyase-Activating Enzyme: Strictly Anaerobic Isolation Yields Active Enzyme Containing a [3Fe–4S]+ Cluster. Biochem. Biophys. Res. Commun. 2000, 269 (2), 451–456. DOI: 10.1006/bbrc.2000.2313.

(44) Cáceres, J. C.; Dolmatch, A.; Greene, B. L. The Mechanism of Inhibition of Pyruvate Formate Lyase by Methacrylate. J. Am. Chem. Soc. 2023, 145 (41), 22504–22515. DOI: 10.1021/jacs.3c07256.

(45) Cáceres, J. C.; Bailey, C. A.; Yokoyama, K.; Greene, B. L. Selenocysteine Substitutions in Thiyl Radical Enzymes. In Methods in Enzymology; Elsevier, 2022; Vol. 662, pp 119–141. DOI: 10.1016/bs.mie.2021.10.014.

(46) Guerrero, F.; Ciragan, A.; Iwaï, H. Tandem SUMO Fusion Vectors for Improving Soluble Protein Expression and Purification. Protein Expr. Purif. 2015, 116, 42–49. DOI: 10.1016/j.pep.2015.08.019.

(47) van den Berg, S.; Löfdahl, P.-Å.; Härd, T.; Berglund, H. Improved Solubility of TEV Protease by Directed Evolution. J. Biotechnol. 2006, 121 (3), 291–298. DOI: 10.1016/j.jbiotec.2005.08.006.

(48) Johnson, D. C.; Unciuleac, M.-C.; Dean, D. R. Controlled Expression and Functional Analysis of Iron-Sulfur Cluster Biosynthetic Components within Azotobacter Vinelandii. J. Bacteriol. 2006, 188 (21), 7551–7561. DOI: 10.1128/JB.00596-06.

(49) Lanz, N. D.; Grove, T. L.; Gogonea, C. B.; Lee, K.-H.; Krebs, C.; Booker, S. J. RlmN and AtsB as Models for the Overproduction and Characterization of Radical SAM Proteins. In Methods in Enzymology; Elsevier, 2012; Vol. 516, pp 125–152. DOI: 10.1016/B978-012-394291-3.00030-7.

(50) Wei, Y.; Mathies, G.; Yokoyama, K.; Chen, J.; Griffin, R. G.; Stubbe, J. A Chemically Competent Thiosulfuranyl Radical on the Escherichia Coli Class III Ribonucleotide Reductase. J. Am. Chem. Soc. 2014, 136 (25), 9001–9013. DOI: 10.1021/ja5030194.

(51) Zhang, L.; King, E.; Luo, R.; Li, H. Development of a High-Throughput, In Vivo Selection Platform for NADPH-Dependent Reactions Based on Redox Balance Principles. ACS Synth. Biol. 2018, 7 (7), 1715–1721. DOI: 10.1021/acssynbio.8b00179.

(52) Gibson, D. G.; Young, L.; Chuang, R.-Y.; Venter, J. C.; Hutchison, C. A.; Smith, H. O. Enzymatic Assembly of DNA Molecules up to Several Hundred Kilobases. Nat. Methods 2009, 6 (5), 343–345. DOI: 10.1038/nmeth.1318.

(53) Bailey, C. A.; Greene, B. L. A Fluorometric Assay for High-Throughput Phosphite Quantitation in Biological and Environmental Matrices. Analyst 2023, 148 (15), 3650–3658. DOI: 10.1039/D3AN00575E.

(54) Edelhoch, H. Spectroscopic Determination of Tryptophan and Tyrosine in Proteins. Biochemistry 1967, 6 (7), 1948–1954. DOI: 10.1021/bi00859a010.

(55) Pace, C. N.; Vajdos, F.; Fee, L.; Grimsley, G.; Gray, T. How to Measure and Predict the Molar Absorption Coefficient of a Protein. Protein Sci. 1995, 4 (11), 2411–2423. DOI: 10.1002/pro.5560041120.

(56) Conradt, H.; Hohmann-Berger, M.; Hohmann, H.-P.; Blaschkowski, H. P.; Knappe, J. Pyruvate Formate-Lyase (Inactive Form) and Pyruvate Formate-Lyase Activating Enzyme of Escherichia Coli: Isolation and Structural Properties. Arch. Biochem. Biophys. 1984, 228 (1), 133–142. DOI: 10.1016/0003-9861(84)90054-7.

(57) Brush, E. J.; Lipsett, K. A.; Kozarich, J. W. Inactivation of Escherichia Coli Pyruvate Formate-Lyase by Hypophosphite: Evidence for a Rate-Limiting Phosphorus-Hydrogen Bond Cleavage. Biochemistry 1988, 27 (6), 2217–2222. DOI: 10.1021/bi00406a061.

(58) Byer, A. S.; McDaniel, E. C.; Impano, S.; Broderick, W. E.; Broderick, J. B. Mechanistic Studies of Radical SAM Enzymes: Pyruvate Formate-Lyase Activating Enzyme and Lysine 2,3-Aminomutase Case Studies. In Methods in Enzymology; Elsevier, 2018; Vol. 606, pp 269–318. DOI: 10.1016/bs.mie.2018.04.013.

(59) Horecker, B. L.; Kornberg, A. The Extinction Coefficients of the Reduced Band of Pyridine Nucleotides. J. Biol. Chem. 1948, 175 (1), 385–390. DOI: 10.1016/S0021-9258(18)57268-9.

(60) Lin, R.; Farmer, P. J. The HNO Adduct of Myoglobin: Synthesis and Characterization. J. Am. Chem. Soc. 2000, 122 (10), 2393– 2394. DOI: 10.1021/ja994079n.

(61) Tofani, L.; Feis, A.; Snoke, R. E.; Berti, D.; Baglioni, P.; Smulevich, G. Spectroscopic and Interfacial Properties of Myoglobin/Surfactant Complexes. Biophys. J. 2004, 87 (2), 1186–1195. DOI: 10.1529/biophysj.104.041731.

(62) Kushida, T.; Ahn, J. S.; Hirata, K.; Kurita, A. Glass Transition of Deoxymyoglobin Probed by Optical Absorption Spectroscopy. Biochem. Biophys. Res. Commun. 1989, 160 (2), 948–953. DOI: 10.1016/0006-291X(89)92527-8.

(63) Kamarei, A. R.; Karel, M. An Improved Method for Preparation of Nitric Oxide Myoglobin. J. Food Sci. 1982, 47 (2), 682–683. DOI: 10.1111/j.1365-2621.1982.tb10155.x.

(64) Unkrig, V.; Neugebauer, F. A.; Knappe, J. The Free Radical of Pyruvate Formate-Lyase. Eur. J. Biochem. 1989, 184 (3), 723–728. DOI: 10.1111/j.1432-1033.1989.tb15072.x.

(65) Stoll, S.; Schweiger, A. EasySpin, a Comprehensive Software Package for Spectral Simulation and Analysis in EPR. J. Magn. Reson. 2006, 178 (1), 42–55. DOI: 10.1016/j.jmr.2005.08.013.

(66) MacLean, B.; Tomazela, D. M.; Shulman, N.; Chambers, M.; Finney, G. L.; Frewen, B.; Kern, R.; Tabb, D. L.; Liebler, D. C.; MacCoss, M. J. Skyline: An Open Source Document Editor for Creating and Analyzing Targeted Proteomics Experiments. Bioinformatics 2010, 26 (7), 966–968. DOI: 10.1093/bioinformatics/btq054.

(67) Pino, L. K.; Searle, B. C.; Bollinger, J. G.; Nunn, B.; MacLean, B.; MacCoss, M. J. The Skyline Ecosystem: Informatics for Quantitative Mass Spectrometry Proteomics. Mass Spectrom. Rev. 2020, 39 (3), 229–244. DOI: 10.1002/mas.21540.

(68) Keefer, L. K.; Nims, R. W.; Davies, K. M.; Wink, D. A. “NONOates” (1-Substituted Diazen-1-Ium-1,2-Diolates) as Nitric Oxide Donors: Convenient Nitric Oxide Dosage Forms. In Methods in Enzymology; Nitric Oxide Part A: Sources and Detection of NO; NO Synthase; Academic Press, 1996; Vol. 268, pp 281–293. DOI: 10.1016/S0076-6879(96)68030-6.

(69) Maragos, C. M.; Morley, D.; Wink, D. A.; Dunams, T. M.; Saavedra, J. E.; Hoffman, A.; Bove, A. A.; Isaac, L.; Hrabie, J. A.; Keefer, L. K. Complexes of .NO with Nucleophiles as Agents for the Controlled Biological Release of Nitric Oxide. Vasorelaxant Effects. J. Med. Chem. 1991, 34 (11), 3242–3247. DOI: 10.1021/jm00115a013.

(70) Moore, E. G.; Gibson, Q. H. Cooperativity in the Dissociation of Nitric Oxide from Hemoglobin. J. Biol. Chem. 1976, 251 (9), 2788–2794. DOI: 10.1016/S0021-9258(17)33557-3.

(71) Møller, J. K. S.; Skibsted, L. H. Nitric Oxide and Myoglobins. Chem. Rev. 2002, 102 (4), 1167–1178. DOI: 10.1021/cr000078y.

(72) Andorfer, M. C.; Backman, L. R. F.; Li, P. L.; Ulrich, E. C.; Drennan, C. L. Rescuing Activity of Oxygen-Damaged Pyruvate Formate-Lyase by a Spare Part Protein. J. Biol. Chem. 2021, 297 (6), 101423. DOI: 10.1016/j.jbc.2021.101423.

(73) Gaston, B. Nitric Oxide and Thiol Groups. Biochim. Biophys. Acta, Bioenerg. 1999, 1411 (2), 323–333. DOI: 10.1016/S0005-2728(99)00023-7.

(74) Madej, E.; Folkes, L. K.; Wardman, P.; Czapski, G.; Goldstein, S. Thiyl Radicals React with Nitric Oxide to Form S-Nitrosothiols with Rate Constants near the Diffusion-Controlled Limit. Free Radic. Biol. Med. 2008, 44 (12), 2013–2018. DOI: 10.1016/j.freeradbiomed.2008.02.015.

(75) Rissanen, M. P.; Ihlenborg, M.; Pekkanen, T. T.; Timonen, R. S. Kinetics of Several Oxygen-Containing Carbon-Centered Free Radical Reactions with Nitric Oxide. J. Phys. Chem. A 2015, 119 (28), 7734–7741. DOI: 10.1021/acs.jpca.5b01027.

(76) Foster, M. W.; Cowan, J. A. Chemistry of Nitric Oxide with Protein-Bound Iron Sulfur Centers. Insights on Physiological Reactivity. J. Am. Chem. Soc. 1999, 121 (17), 4093–4100. DOI: 10.1021/ja9901056.

(77) Vanin, A. F. Research into Dinitrosyl Iron Complexes in Living Organisms Through EPR as an Example of Applying This Method in Biology: A Review. Appl. Magn. Reson. 2023, 54 (2), 289– 309. DOI: 10.1007/s00723-022-01518-3.

(78) Lehnert, N.; Kim, E.; Dong, H. T.; Harland, J. B.; Hunt, A. P.; Manickas, E. C.; Oakley, K. M.; Pham, J.; Reed, G. C.; Alfaro, V. S. The Biologically Relevant Coordination Chemistry of Iron and Nitric Oxide: Electronic Structure and Reactivity. Chem. Rev. 2021, 121 (24), 14682–14905. DOI: 10.1021/acs.chemrev.1c00253.

(79) Tinberg, C. E.; Tonzetich, Z. J.; Wang, H.; Do, L. H.; Yoda, Y.; Cramer, S. P.; Lippard, S. J. Characterization of Iron Dinitrosyl Species Formed in the Reaction of Nitric Oxide with a Biological Rieske Center. J. Am. Chem. Soc. 2010, 132 (51), 18168–18176. DOI: 10.1021/ja106290p.

(80) Tsou, C.-C.; Lu, T.-T.; Liaw, W.-F. EPR, UV-Vis, IR, and X-Ray Demonstration of the Anionic Dimeric Dinitrosyl Iron Complex [(NO)2Fe(μ-StBu)2Fe(NO)2]?: Relevance to the Products of Nitrosylation of Cytosolic and Mitochondrial Aconitases, and High-Potential Iron Proteins. J. Am. Chem. Soc. 2007, 129 (42), 12626– 12627. DOI: 10.1021/ja0751375.

(81) Cruz-Ramos, H.; Crack, J.; Wu, G.; Hughes, M. N.; Scott, C.; Thomson, A. J.; Green, J.; Poole, R. K. NO Sensing by FNR: Regulation of the Escherichia Coli NO-Detoxifying Flavohaemoglobin, Hmp. EMBO J. 2002, 21 (13), 3235–3244. DOI: 10.1093/emboj/cdf339.

(82) Ma, K.; Xue, B.; Chu, R.; Zheng, Y.; Sharma, S.; Jiang, L.; Hu, M.; Xie, Y.; Hu, Y.; Tao, T.; Zhou, Y.; Liu, D.; Li, Z.; Yang, Q.; Chen, Y.; Wu, S.; Tong, Y.; Robinson, R. C.; Yew, W. S.; Jin, X.; Liu, Y.; Zhao, H.; Ang, E. L.; Wei, Y.; Zhang, Y. A Widespread Radical-Mediated Glycolysis Pathway. J. Am. Chem. Soc. 2024, 146 (38), 26187–26197. DOI: 10.1021/jacs.4c07718.

(83) Greene, B. L.; Kang, G.; Cui, C.; Bennati, M.; Nocera, D. G.; Drennan, C. L.; Stubbe, J. Ribonucleotide Reductases: Structure, Chemistry, and Metabolism Suggest New Therapeutic Targets. Annu. Rev. Biochem. 2020, 89 (1), 45–75. DOI: 10.1146/annurevbiochem-013118-111843.

(84) Mulliez, E.; Fontecave, M.; Gaillard, J.; Reichard, P. An Iron-Sulfur Center and a Free Radical in the Active Anaerobic Ribonucleotide Reductase of Escherichia Coli. J. Biol. Chem. 1993, 268 (4), 2296–2299. DOI: 10.1016/S0021-9258(18)53772-8.

(85) Yokoyama, K.; Lilla, E. A. C–C Bond Forming Radical SAM Enzymes Involved in the Construction of Carbon Skeletons of Cofactors and Natural Products. Nat. Prod. Rep. 2018, 35 (7), 660– 694. DOI: 10.1039/c8np00006a.

(86) Lilla, E. A.; Yokoyama, K. Carbon Extension in Peptidylnucleoside Biosynthesis by Radical SAM Enzymes. Nat. Chem. Biol. 2016, 12 (11), 905–907. DOI: 10.1038/nchembio.2187.

(87) Bogdan, C. Nitric Oxide Synthase in Innate and Adaptive Immunity: An Update. Trends Immunol. 2015, 36 (3), 161–178. DOI: 10.1016/j.it.2015.01.003.

(88) Bogdan, C. Reactive Oxygen and Reactive Nitrogen Intermediates in the Immune System. In The Immune Response to Infection; Kaufmann, S. H. E., Rouse, B. T., Sacks, D. L., Eds.; ASM Press: Washington, DC, USA, 2014; pp 69–84. DOI: 10.1128/9781555816872.ch5.

(89) Bogdan, C. Nitric Oxide and the Immune Response. Nat. Immunol. 2001, 2 (10), 907–916. DOI: 10.1038/ni1001-907.

(90) Tsai, W. C.; Strieter, R. M.; Zisman, D. A.; Wilkowski, J. M.; Bucknell, K. A.; Chen, G. H.; Standiford, T. J. Nitric Oxide Is Required for Effective Innate Immunity Against Klebsiella Pneumoniae. Infect. Immun. 1997, 65 (5), 1870–1875. DOI: 10.1128/iai.65.5.1870-1875.1997.

(91) Alam, M. S.; Akaike, T.; Okamoto, S.; Kubota, T.; Yoshitake, J.; Sawa, T.; Miyamoto, Y.; Tamura, F.; Maeda, H. Role of Nitric Oxide in Host Defense in Murine Salmonellosis as a Function of Its Antibacterial and Antiapoptotic Activities. Infect. Immun. 2002, 70 (6), 3130–3142. DOI: 10.1128/IAI.70.6.3130-3142.2002.

(92) Hall, J. R.; Rouillard, K. R.; Suchyta, D. J.; Brown, M. D.; Ahonen, M. J. R.; Schoenfisch, M. H. Mode of Nitric Oxide Delivery Affects Antibacterial Action. ACS Biomater. Sci. Eng. 2020, 6 (1), 433– 441. DOI: 10.1021/acsbiomaterials.9b01384.

(93) Privett, B. J.; Broadnax, A. D.; Bauman, S. J.; Riccio, D. A.; Schoenfisch, M. H. Examination of Bacterial Resistance to Exogenous Nitric Oxide. Nitric Oxide 2012, 26 (3), 169–173. DOI: 10.1016/j.niox.2012.02.002.

(94) Schapiro, J. M.; Libby, S. J.; Fang, F. C. Inhibition of Bacterial DNA Replication by Zinc Mobilization During Nitrosative Stress. Proc. Natl. Acad. Sci. U.S.A. 2003, 100 (14), 8496–8501. DOI: 10.1073/pnas.1033133100.

(95) Lepoivre, M.; Fieschi, F.; Coves, J.; Thelander, L.; Fontecave, M. Inactivation of Ribonucleotide Reductase by Nitric Oxide. Biochem. Biophys. Res. Commun. 1991, 179 (1), 442–448. DOI: 10.1016/0006-291X(91)91390-X.

(96) Roy, B.; Lepoivre, M.; Henry, Y.; Fontecave, M. Inhibition of Ribonucleotide Reductase by Nitric Oxide Derived from Thio-nitrites: Reversible Modifications of Both Subunits. Biochemistry 1995, 34 (16), 5411–5418. DOI: 10.1021/bi00016a012.

(97) Butler, C. S.; Seward, H. E.; Greenwood, C.; Thomson, A. J. Fast Cytochrome Bo from Escherichia Coli Binds Two Molecules of Nitric Oxide at CuB. Biochemistry 1997, 36 (51), 16259–16266. DOI: 10.1021/bi971481a.

(98) Richardson, A. R.; Payne, E. C.; Younger, N.; Karlinsey, J. E.; Thomas, V. C.; Becker, L. A.; Navarre, W. W.; Castor, M. E.; Libby, S. J.; Fang, F. C. Multiple Targets of Nitric Oxide in the Tricarboxylic Acid Cycle of Salmonella Enterica Serovar Typhimurium. Cell Host Microbe 2011, 10 (1), 33–43. DOI: 10.1016/j.chom.2011.06.004.

(99) Schairer, D. O.; Chouake, J. S.; Nosanchuk, J. D.; Friedman, A. J. The Potential of Nitric Oxide Releasing Therapies as Antimicrobial Agents. Virulence 2012, 3 (3), 271–279. DOI: 10.4161/viru.20328.

(100) Parast, C. V.; Wong, K. K.; Lewisch, S. A.; Kozarich, J. W.; Peisach, J.; Magliozzo, R. S. Hydrogen Exchange of the Glycyl Radical of Pyruvate Formate-Lyase Is Catalyzed by Cysteine 419. Biochemistry 1995, 34 (8), 2393–2399. DOI: 10.1021/bi00008a001.

(101) Bartberger, M. D.; Mannion, J. D.; Powell, S. C.; Stamler, J. S.; Houk, K. N.; Toone, E. J. S-N Dissociation Energies of S-Nitrosothiols: On the Origins of Nitrosothiol Decomposition Rates. J. Am. Chem. Soc. 2001, 123 (36), 8868–8869. DOI: 10.1021/ja0109390.

(102) Stamler, J. S.; Toone, E. J. The Decomposition of Thio-nitrites. Curr. Opin. Chem. Biol. 2002, 6 (6), 779–785. DOI: 10.1016/S1367-5931(02)00383-6.

(103) Forrester, M. T.; Foster, M. W.; Benhar, M.; Stamler, J. S. Detection of Protein S-Nitrosylation with the Biotin-Switch Technique. Free Radic. Biol. Med. 2009, 46 (2), 119–126. DOI: 10.1016/j.freeradbiomed.2008.09.034.

(104) Leclerc, M.; Bedu-Ferrari, C.; Etienne-Mesmin, L.; Mariadassou, M.; Lebreuilly, L.; Tran, S.-L.; Brazeau, L.; Mayeur, C.; Delmas, J.; Rué, O.; Denis, S.; Blanquet-Diot, S.; Ramarao, N. Nitric Oxide Impacts Human Gut Microbiota Diversity and Functionalities. mSystems 2021, 6 (5), e00558–21. DOI: 10.1128/msystems.00558-21.

(105) Wang, Q.; Ou, M. S.; Kim, Y.; Ingram, L. O.; Shanmugam, K. T. Metabolic Flux Control at the Pyruvate Node in an Anaerobic Escherichia Coli Strain with an Active Pyruvate Dehydrogenase. Appl. Environ. Microbiol. 2010, 76 (7), 2107–2114. DOI: 10.1128/AEM.02545-09.

(106) Kaiser, M.; Sawers, G. Pyruvate Formate-Lyase Is Not Essential for Nitrate Respiration by Escherichia Coli. FEMS Microbiol. Lett. 1994, 117 (2), 163–168. DOI: 10.1111/j.1574-6968.1994.tb06759.x.

(107) Duan, Y.; Wei, Y.; Xing, M.; Liu, J.; Jiang, L.; Lu, Q.; Liu, X.; Liu, Y.; Ang, E. L.; Liao, R.-Z.; Yuchi, Z.; Zhao, H.; Zhang, Y. Anaerobic Hydroxyproline Degradation Involving C–N Cleavage by a Glycyl Radical Enzyme. J. Am. Chem. Soc. 2022, 144 (22), 9715–9722. DOI: 10.1021/jacs.2c01673.

(108) Craciun, S.; Balskus, E. P. Microbial Conversion of Choline to Trimethylamine Requires a Glycyl Radical Enzyme. Proc. Natl. Acad. Sci. U.S.A. 2012, 109 (52), 21307–21312. DOI: 10.1073/pnas.1215689109.

(109) Dawson, C. D.; Irwin, S. M.; Backman, L. R. F.; Le, C.; Wang, J. X.; Vennelakanti, V.; Yang, Z.; Kulik, H. J.; Drennan, C. L.; Balskus, E. P. Molecular Basis of C–S Bond Cleavage in the Glycyl Radical Enzyme Isethionate Sulfite-Lyase. Cell Chem. Biol. 2021, 28 (9), 1333– 1346.e7. DOI: 10.1016/j.chembiol.2021.03.001.

(110) Liu, D.; Wei, Y.; Liu, X.; Zhou, Y.; Jiang, L.; Yin, J.; Wang, F.; Hu, Y.; Nanjaraj Urs, A. N.; Liu, Y.; Ang, E. L.; Zhao, S.; Zhao, H.; Zhang, Y. Indoleacetate Decarboxylase Is a Glycyl Radical Enzyme Catalyzing the Formation of Malodorant Skatole. Nat. Commun. 2018, 9 (1), 4224. DOI: 10.1038/s41467-018-06627-x.

(111) Levin, B. J.; Balskus, E. P. Discovering Radical-Dependent Enzymes in the Human Gut Microbiota. Curr. Opin. Chem. Biol. 2018, 47, 86–93. DOI: 10.1016/j.cbpa.2018.09.011.

(112) Uyeda, K.; Rabinowitz, J. C. Pyruvate-Ferredoxin Oxidoreductase: III. PURIFICATION AND PROPERTIES OF THE ENZYME. J. Biol. Chem. 1971, 246 (10), 3111–3119. DOI: 10.1016/S0021-9258(18)62202-1

