## Supplemental Information for "Nitric Oxide Inhibition of Glycyl Radical Enzymes and Their Activases"

**Table S1. List of Primers**

| Plasmid name | Primer name | Sequence |
|---|---|---|
| pHYRSF53-PFL-AE | Fragment forward | 5′-tggatcccatatgtcagttattggtcgcattcact-3′ |
| | Fragment reverse | 5′-agcctaggttaattagaacattaccttatgaccgtactgct-3′ |
| | Vector forward | 5′-taatgttctaattaacctaggctgctgccacc-3′ |
| | Vector reverse | 5′-taactgacatatgggatccaccaatctgttctctgt-3′ |
| pHYRSF53-FDH | Fragment forward | 5′-tggatccatggcgaaggttttgtgcg-3′ |
| | Fragment reverse | 5′-taggttaattaaacggctttcttaaacttcgcag-3′ |
| | Vector forward | 5′-agaaagccgtttaattaacctaggctgctgccacc-3′ |
| | Vector reverse | 5′-ttcgccatggatccaccaatctgttctctgtgag-3′ |
| pHYRSF53-NrdG | Fragment forward | 5′-gctcacagaacagattggtgatcccatatgaattatcatcagtactatcc-3′ |
| | Fragment reverse | 5′-ggtggcagcagcctaggttaatcatcgcaaatgatgcacc-3′ |
| | Vector forward | 5′-ggtgcatcatttgcgatgattaacctaggctgctgccaccgc-3′ |
| | Vector reverse | 5′-gtactgatgataattcatatgggatccaccaatctgttctctg-3′ |

### Supplemental Methods

*Protein Expression and Purification.*

For the expression of SUMO-FDH, *E. coli* BL21(DE3) cells transformed with the corresponding plasmid were inoculated into a 15 mL preculture of LB broth supplemented with 50 μg/mL of kanamycin. The preculture was incubated overnight at 37 °C in a 50 mL centrifuge tube shaking at 200 rpm. The following day, 15 mL of the overnight preculture were inoculated into 1.5 L of LB broth, supplemented with 50 μg/mL of kanamycin, and grown at 37 °C, shaking at 200 rpm. Protein expression was induced by adding 1 mM IPTG when the $OD_{600}$ reached 1.0, then the culture was maintained at 20 °C overnight. Cells were harvested by centrifugation at 8,000 × g for 10 min, and the collected cell paste was flash frozen in liquid $N_2$ and stored at –80 °C until purification.

For FDH purification, we resuspended the cell paste in 5 mL/g cell paste resuspension buffer consisting of 50 mM Tris, 300 mM KCl, 0.5 mM TCEP, and 5% glycerol, adjusted to pH 8.0, and lysed the cells by French press in an Emulsiflex C3 homogenizer at 14,000 psi. Every following step was performed at 4 °C. The cell extract was clarified by centrifugation at 30,000 × g for 30 min and the cell debris was discarded. To remove nucleic acids, 1.5% w/v streptomycin sulfate was added dropwise while stirring and incubated for 10 min before being centrifuged at 30,000 × g for 30 min. The pellet was discarded, and the supernatant was filtered through a 0.65 μm filter. The clarified lysate was applied to a Ni-NTA column equilibrated with wash buffer composed of 25 mM imidazole, 50 mM Tris, 300 mM KCl, 0.5 mM TCEP and 5% glycerol at pH 8.0. The resin was washed with 20 column volumes of wash buffer and eluted with elution buffer in which the imidazole concentration was raised to 200 mM. Fractions containing protein were pooled and concentrated using Amicon Ultra-15 10 kDa molecular weight cut-off (MWCO) centrifugal filter units and then buffer exchanged to resuspension buffer using a HiTrap desalting column. To remove the N-terminal His-tag, SUMO-FDH was incubated with ULP1 protease in a 1:100 ratio of protease to protein overnight and then the digestion product was injected back into the Ni-NTA column to remove the cleaved N-terminal tag, His-tagged protease, and undigested SUMO-FDH. The Ni-NTA flowthrough was collected, and the purified, untagged protein was concentrated and desalted as described above. The purified protein was stored in resuspension buffer supplemented with 20% (w/v) glycerol at –80 °C until used.

SUMO-PFL-AE and SUMO-RNR-AE were expressed in similar conditions. We transformed BL21(DE3) *E. coli* with the respective plasmid and inoculated the cells in a 100 mL preculture of LB broth supplemented with 50 μg/mL of kanamycin. For SUMO-RNR-AE we additionally transformed the cells with the plasmid pDB1282, which encodes for the *isc* operon from *Azotobacter vinelandii* involved in iron-sulfur cluster biosynthesis, and supplemented the cultures with 100 μg/mL of carbenicillin and 0.1% (w/v) of L-arabinose. The preculture was grown overnight at 37 °C in a 250 mL flask shaking at 200 rpm. The following day, 10 mL of the overnight preculture were inoculated into 2 L of terrific broth, supplemented with 50 μg/mL of kanamycin, and grown at 37 °C with shaking at 200 rpm. Expression was induced by adding 0.25 mM IPTG at an $OD_{600}$ of 1.0, and the culture was supplemented with 0.5 mM L-cysteine, and 0.5 mM $(NH_4)_2Fe^{II}(SO_4)_2$, before lowering the temperature to 30 °C and shaking for another 4 h at 200 rpm. Next, L-cysteine and $(NH_4)_2Fe^{II}(SO_4)_2$ were added to a final concentration of 1 mM and the cultures were bubbled with argon for 1 h at 20 °C before being sealed with a rubber plug and incubated at 20 °C and 100 rpm overnight. Finally, 1 mM TCEP was added to the culture and the cells were harvested by centrifugation at 8,000 × g for 10 min. The collected cell paste was flash frozen in liquid $N_2$ and stored at –80 °C until purification.

To purify PFL-AE and RNR-AE we followed the same protocol. The frozen cell paste was moved into a Coy glovebox (<20 ppm $O_2$) and resuspended in “resuspension” buffer consisting of 50 mM potassium phosphate, 150 mM KCl, 1 mM DTT and 5% (w/v) glycerol, adjusted to pH 8.0, to 4 mL/g wet cell paste, and lysed by passage through an Emulsiflex C3 French press homogenizer at 14,000 psi. The sample was briefly exposed to air in this step but was quickly degassed in a Schleck line before continuing the purification. After the lysis step, the following steps were performed in a glovebox or in degassed O-ring sealed centrifuge bottles. The cell extract was clarified by centrifugation at 30,000 × g for 90 min and the cell debris was discarded. The clarified lysate was applied to a HisPur™ Cobalt column equilibrated with “wash” buffer composed of 20 mM imidazole, 50 mM potassium phosphate, 150 mM KCl, 1 mM DTT and 5% (v/v) glycerol at pH 8.0. The resin was washed with 20 column volumes of wash buffer and eluted with “elution” buffer, in which the imidazole concentration was raised to 200 mM. Brown-colored fractions were pooled and concentrated using a Vivaspin 20 filtration units of 10 kDa MWCO. The concentrated protein was then diluted to <20 mM imidazole in resuspension buffer. To remove the N-terminal poly-histidine tag attached to the SUMO tag, SUMO-PFL-AE or SUMO-RNR-AE were incubated with ULP1 in a 1:100 ratio of protease to protein for 1 h and then the digestion product was applied onto the HisPur™ Cobalt column to remove the cleaved N-terminal tag, his-tagged ULP1 protease, and undigested SUMO-PFL-AE. The HisPur™ Cobalt flow-through was collected and the untagged PFL-AE protein was concentrated in 10 kDa MWCO Vivaspin 20 filtration units and then buffer exchanged to resuspension buffer using a HiTrap desalting column and stored in resuspension buffer supplemented with 20% (w/v) glycerol at -80 °C until used.

We recombinantly expressed the *E. coli* class III RNR via transformation of the plasmid pET28a-EcNrdD into BL21 (DE3) *E. coli* cells and plated transformants on 50 µg/mL kanamycin LB agar plates. A single colony was then picked and inoculated in a 15 mL preculture of LB broth in a 50 mL falcon tube, supplemented with 50 µg/mL of kanamycin and incubated the culture overnight at 37 °C shaking at 200 rpm. Next, 15 mL of saturated overnight cultures were used to inoculate 1.5 L of LB, supplemented with 50 µg/mL of kanamycin and 50 µM zinc acetate, and grown at 37 °C at 180 rpm. Once an $OD_{600}$ ~0.6 was reached, protein expression was induced by adding 1 mM ITPG and the cells were maintained at 37 °C for 4 h. After 4 h, the cells were pelleted by centrifugation 3,000 × g for 10 min at 4 °C and flash frozen in liquid $N_2$ and stored at –80 °C.

To purify RNR we resuspended the cell paste in 5 mL/g cell paste of "resuspension" buffer consisting on 50 mM Tris, 50 mM KCl, 10% (w/v) glycerol, and 1 mM PMSF adjusted to pH 7.6 and homogenized by a glass homogenizer on ice. The homogenized cell suspension was then lysed via sonication, again on ice, using the following settings: an amplitude of 60%, pulse time of 20 s-on and 10 s-off for 6 min per every 45 mL of cell suspension. All following steps were carried out in a cold room at 4 °C. To remove nucleic acids, 1.5% (w/v) streptomycin sulfate was added dropwise while stirring and incubated for 15 min before being centrifuged at 30,000 × g for 30 min. The lysed cell suspension was then further clarified by centrifugation at 30,000 × g for 30 min, and the clarified lysate was then loaded onto a Ni-NTA column, pre-equilibrated with "wash" buffer composed of 40 mM imidazole, 50 mM Tris, and 50 mM KCl adjusted to pH 7.6. The column was washed with 20 column volumes of wash buffer and then the protein was eluted with "elution" buffer containing 250 mM imidazole, 50 mM Tris, and 50 mM KCl adjusted to pH 7.6. Protein-containing fractions were pooled and concentrated using Amicon Ultra-15 of 50 kDa MWCO centrifugal filter units then buffer exchanged to resuspension buffer using a HiTrap desalting column. The buffer-exchanged protein was then supplemented with 10% (w/v) glycerol, flash-frozen in liquid $N_2$, and stored at –80 °C until used.

*Enzymatic Determination of Fermentation Products.*

We quantified the concentrations of the metabolites formate, lactate, and ethanol in anaerobic bacterial culture extracellular media by using enzymatic-coupled assays. These assays employed the enzymes formate dehydrogenase (FDH), lactate dehydrogenase (LDH), and alcohol dehydrogenase (ADH), respectively, to couple the oxidation of the target compounds with the reduction of $NAD^+$ to NADH. For the case of acetate, we used the acetic acid assay kit from Megazyme that couples the acetyl-CoA synthase (ACS) dependent formation of acetyl CoA to the production of NADH by the oxidation of malate to oxaloacetate and condensation to citrate by malate dehydrogenase (MDH) and citrate synthase (CS), respectively.

The assays were conducted in Corning™ Half-area 96-well Clear Flat Bottom UV-Transparent Microplates. Each reaction mixture consisted of 50 μL of sample, 5 mM $NAD^+$, and 400 mM Tris buffer at pH 8.1, with the corresponding enzyme (350 nM LDH, 350 nM ADH, or 1 μM FDH; ACS, MDH and CS concentration as indicate by manufacturer) added to achieve a final volume of 150 μL. Reactions were initiated by adding 100 μL of enzyme-containing reaction mix, incubated at 30 °C for 60 min, and then the formation of NADH was measuring by the UV absorbance at 340 nm using a Tecan Spark plate reader.

Calibration curves were generated using authentic sodium formate, sodium lactate, ethanol, or sodium acetate in water, which were processed identically to the cell media samples. For formate and acetate, the calibration curves demonstrated a linear response in the range of 0 to 15 mM and 0 to 1.7 mM respectively, which was fitted to a linear model for determination of formate/acetate concentrations.

For lactate and ethanol, due to substrate inhibition effects on the enzymes,[1,2] the calibration curves were non-linear in the range of 0 to 30 mM. These were best fitted using a two-phase association model. The two-phase association was modeled using the following system of equations:

$$SpanFast = (Plateau - Y_0) * PercentFast * .01$$

$$SpanSlow = (Plateau - Y_0) * (100 - PercentFast) * .01$$

$$Y = Y_0 + SpanFast * (1 - e^{-KFast*X}) + SpanSlow * (1 - e^{-KSlow*X})$$

Where $Y_0$ is the absorbance at 340 nm when the metabolite concentration (X) is zero, Plateau represents the absorbance at 340 nm at an infinite metabolite concentration. $KFast$ and $KSlow$ are the slopes for the fast and slow phases, respectively and PercentFast represents the fraction of the total span (from $Y_0$ to Plateau) attributable to the faster kinetic component.

To determine the metabolite concentration for each measured absorbance value, we employed numerical methods using the "fsolve" function from the Optimization Toolbox v24.2 in MATLAB Version 24.2.0.2712019 (R2024b). This allowed us to solve for X based on the absorbance values for each sample.

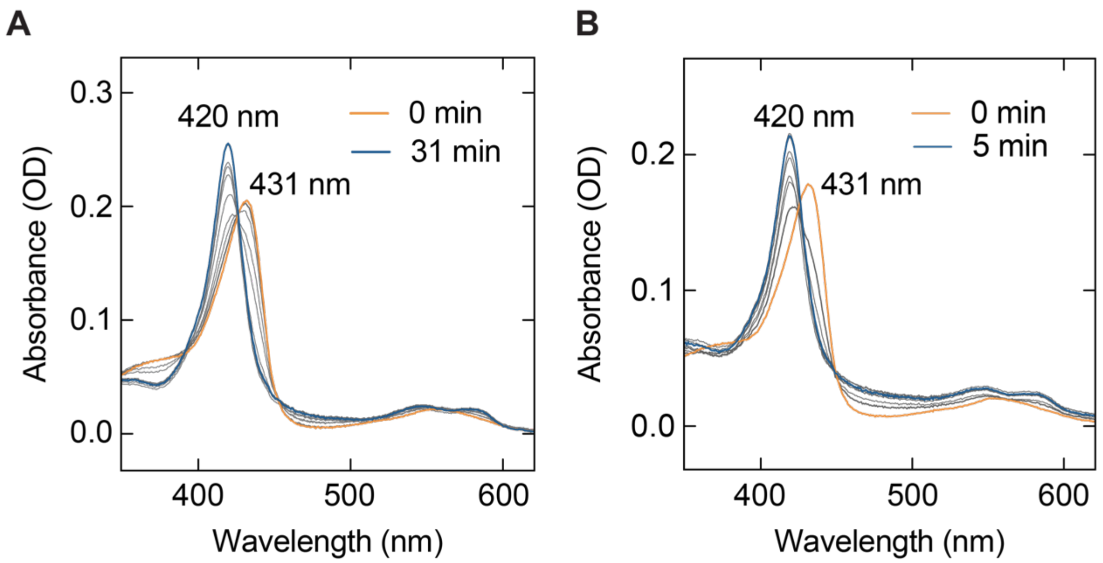

**Figure S1.** UV-vis absorption spectroscopy of NO binding to deoxyMb during the release of NO from NONOate. Samples were prepared identically to those in the aPFL inhibition experiments reported in in the main text **Figure 1**, and NO release was quenched by dilution into buffer consisting of 100 mM glycine at pH 10.0. **A** Reaction of deoxyMb with 100 μM DEA NONOate. **B** Reaction of deoxyMb with 1 mM DEA NONOate. The heme Soret band shifts from 431 nm for the deoxyMb form to 420 nm in the NO-Mb form.

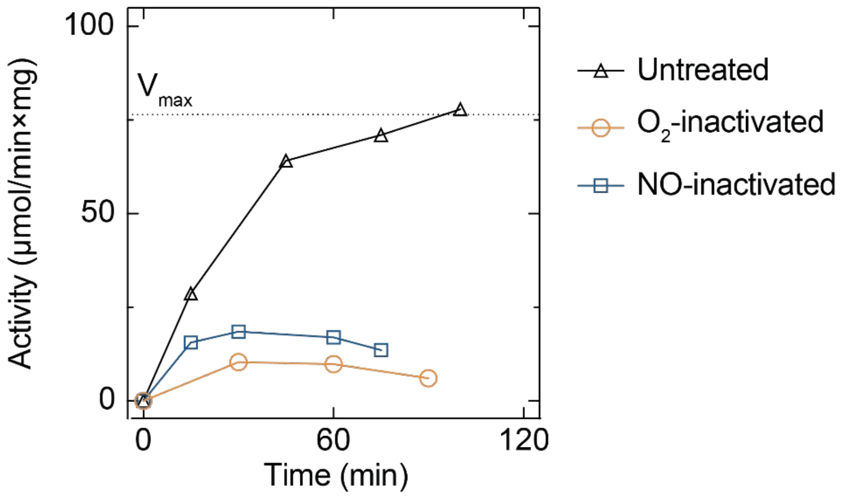

**Figure S2.** Reactivation of PFL inactivated by NO or $O_2$. Reactivation of untreated (black triangles), $O_2$- (orange circles), and NO-inactivated (blue squares) PFL by PFL-AE. PFL was activated with 0.1 molar equivalents of PFL-AE and the activity was measured by the enzyme coupled assay as described in the main text. The reported $V_{max}$ (dashed line) indicates the maximum PFL activation routinely achieved from fresh PFL, as previously determined.[3]

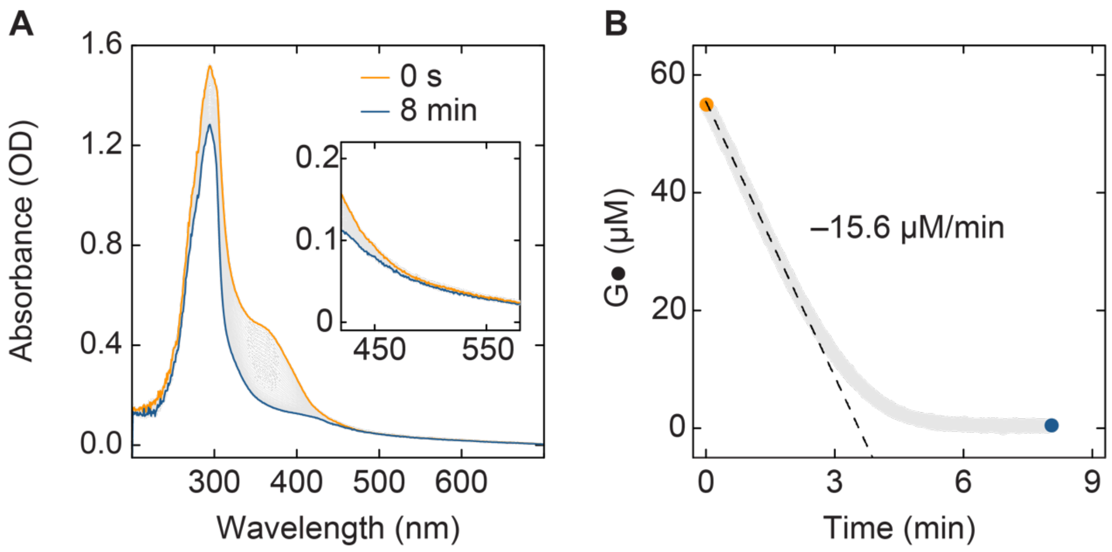

**Figure S3.** Inactivation of aPFL by NO followed by UV-vis absorption spectroscopy. **A** The UV/vis spectrum of 55 μM aPFL during inhibition by NO following the addition of 500 μM NONOate at room temperature and pH 7.6 and incubated for 8 min. No bands between 450 and 550 nm (inset) were observed that could indicate the formation of S-nitrosothiols. **B** The calculated glycyl radical (G●) concentration over the course of the inhibition, estimated by the absorbance at 365 nm and using the reported glycyl radical extinction coefficient of 8,000 $M^{-1}cm^{-1}$.[4]

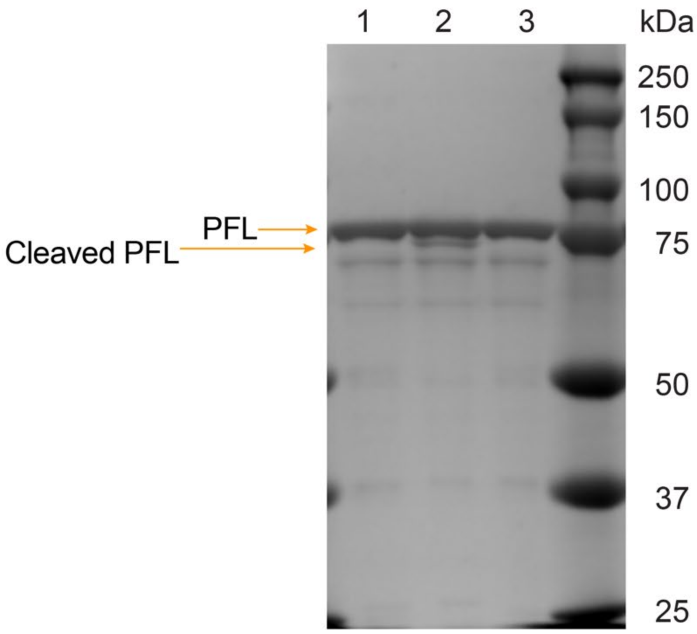

**Figure S4.** Inactivation of aPFL by NO followed by SDS-PAGE. 25 μM aPFL was treated with 100 mM NO for 20 minutes and the sample was then resolved on an 8% bisacrylamide gel (lane 1). Similar samples of aPFL exposed to $O_2$ (lane 2) and non-activated PFL (lane 3) were resolved for comparison.

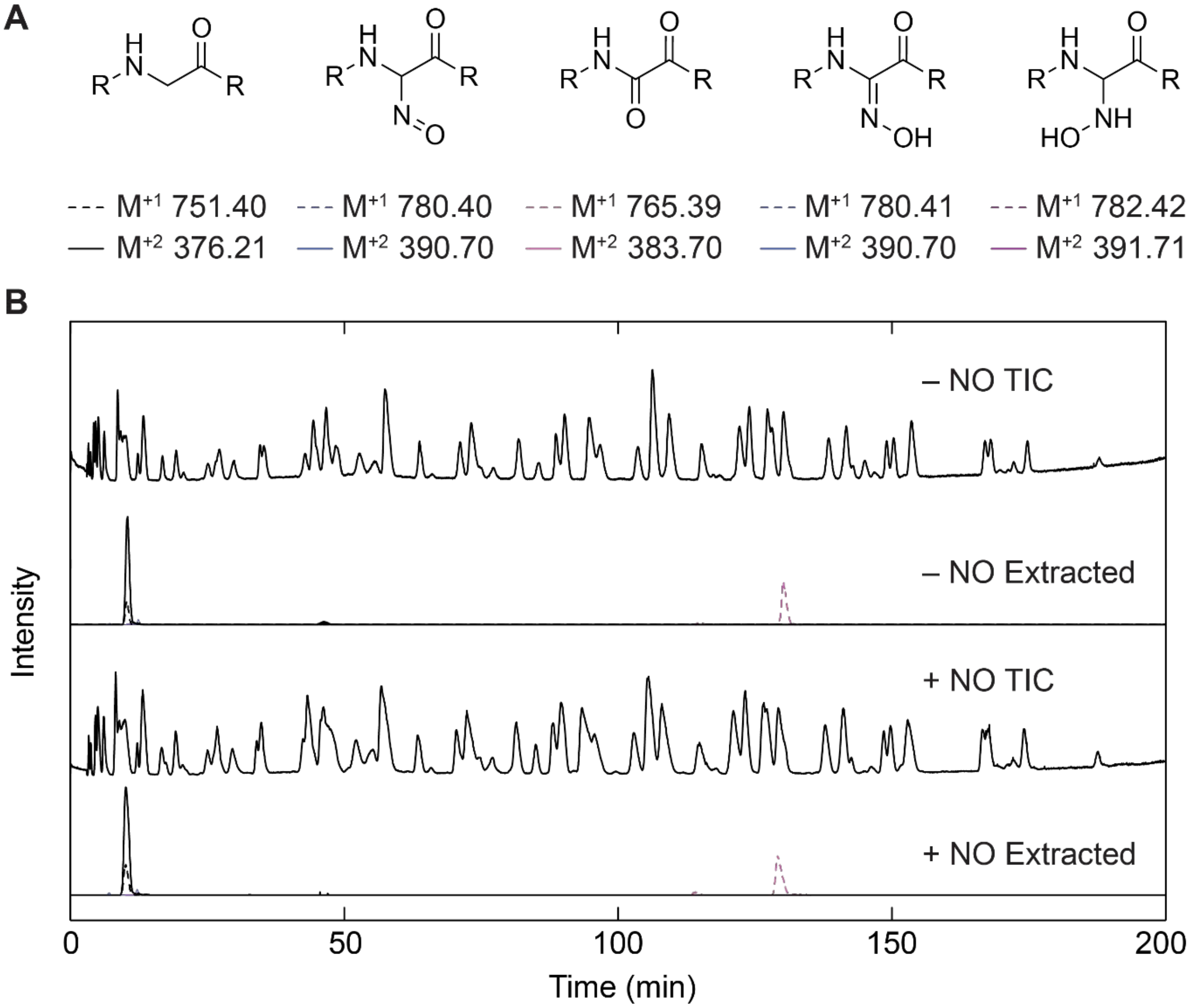

**Figure S5.** LC-MS/MS analysis of tryptic peptides for potential NO-induced post-translational modifications of PFL. **A** Proposed post-translational modifications induced by NO following reaction of the glycyl radical with NO, including alkyl nitroso/oxime, $\alpha$-keto acid, and hydroxylamine. The predicted tryptic digest peptide m/z values are reported. **B** (upper panel) Total ion chromatogram (TIC, top) and extracted mass chromatogram (bottom) for aPFL not treated with NONOate. (lower panel) TIC (top) and extracted mass chromatogram (bottom) for aPFL treated with 1 mM NONOate. A peak corresponding to the $M^{+1}$ ion of the $\alpha$-keto acid modified peptide, with a retention time of 129.5 min, was detected in both NO-treated and untreated samples. However, MS/MS fragmentation analysis revealed that the detected ion's fragmentation pattern did not match the predicted fragmentation for the $\alpha$-keto acid-modified glycyl radical peptide (data not shown).

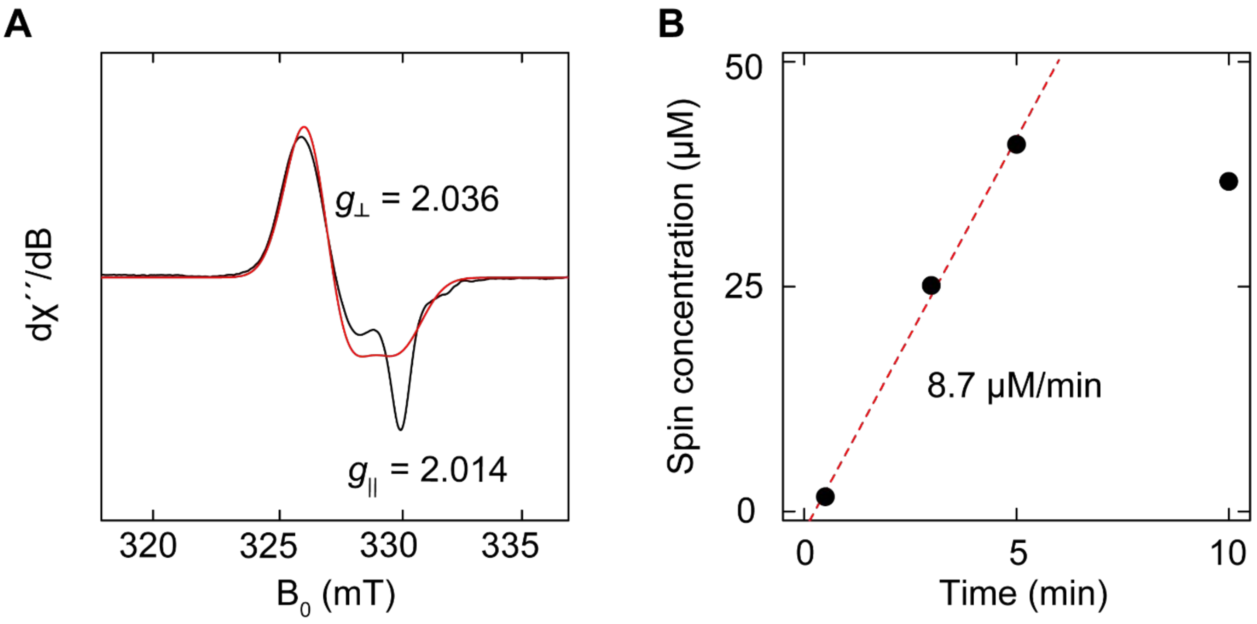

**Figure S6.** X-band EPR spectrum of PFL-AE reacted with NONOate and kinetics. **A** X-band EPR spectra of 25 μM PFL-AE treated with 500 μM NONOate freeze-quenched after 10 min, corresponding to spectra shown in main text **Figure 3** (black) and simulation (red). EPR conditions: microwave frequency, 9.3 GHz; modulation amplitude, 2 G; power 2 mW; temperature, 100 K. 30 scans were averaged. **B** Kinetics of PFL-AE DNIC formation. Spin concentrations were computed from the double integral of the first harmonic signal of the EPR spectra shown in main text **Figure 3** and referenced to a 4-hydroxy-TEMPO standard.

**Table S2. EPR simulation parameters for PFL AE treated with NO.**

| Sample | $g_{\perp}$ | $g_{\parallel}$ | Concentration (μM) | Spin/protein (%) |
|---|---|---|---|---|
| PFL-AE 10 min | 2.00356 (1) | 2.014 (1) | 36.7 | 73 |
| 5 min | | | 40.8 | 82 |
| 3 min | | | 25.2 | 50 |
| 30 s | | | 1.7 | 3 |
| PFL-AE + SAM | 2.0366(1) | 2.0134(3) | 8.7 | 17 |
| Sample | $g_{\perp}$ | $g_{\parallel}$ | Concentration (μM) | Fraction Reduced (%) |
| rPFL-AE | 2.0090 (1) | 1.9703 (1) | 18.9 | 70 |
| rPFL-AE + SAM | 2.00938 (2) | 1.9708 (1) | 25.3 | 75 |

Errors, reported in parenthesis for the last significant digit, are the estimated uncertainty of the EasySpin non-linear least squares fit simulation parameters.

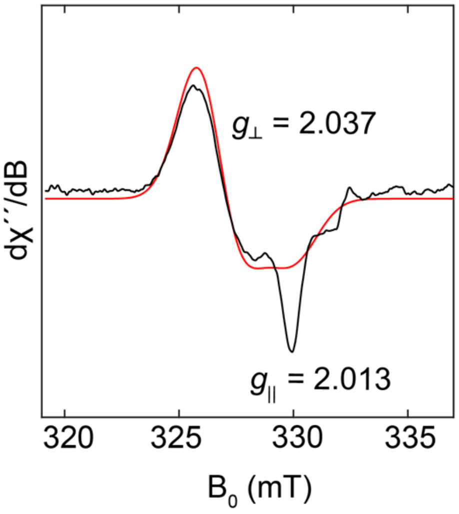

**Figure S7.** X-band EPR spectrum of PFL-AE reacted with NONOate in the presence of SAM. 50 µM PFL-AE was reacted with 500 µM NONOate over 5 min in the presence of 2 mM SAM then frozen in liquid $N_2$-cooled isopentane in an EPR tube. Black trace shows the experimental data and the red trace the simulation. EPR conditions: microwave frequency, 9.3 GHz; modulation amplitude, 2 G; power, 2 mW; temperature, 100 K. 30 scans were averaged.

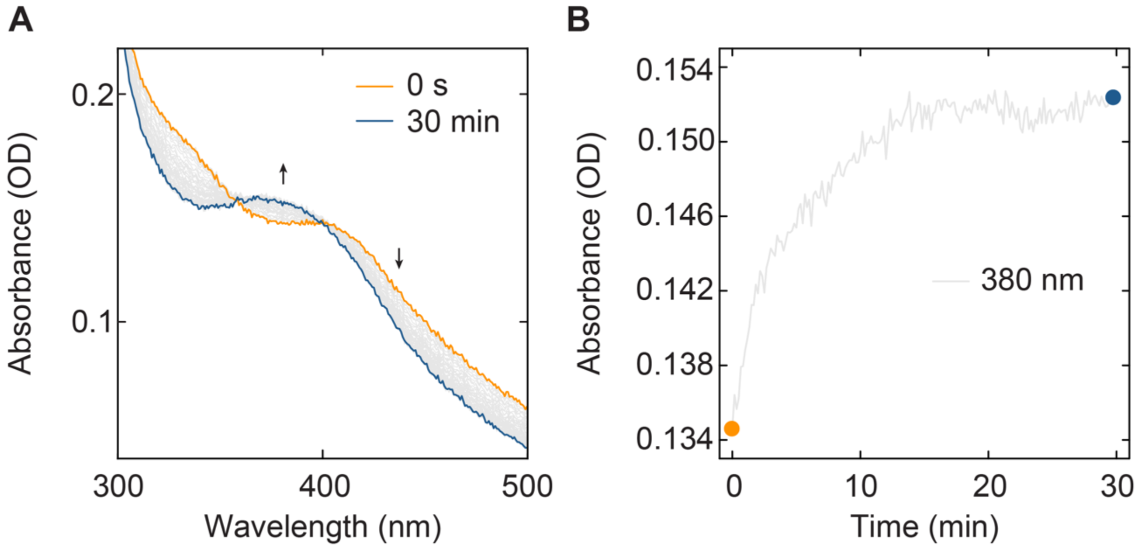

**Figure S8.** UV-vis absorption spectroscopy following inactivation of PFL-AE by NONOate. 20 µM PFL-AE was incubated with 100 µM NONOate at room temperature and pH 7.6 and incubated for 30 min. **A** Time-resolved UV/vis spectra collected every 30 s over 30 min following NONOate addition to PFL-AE from t = 0 (orange) to t = 30 min (blue). **B** Extracted absorption at 380 nm corresponding the production of DNIC complexes from t = 0 min (orange) to t = 30 min (blue).

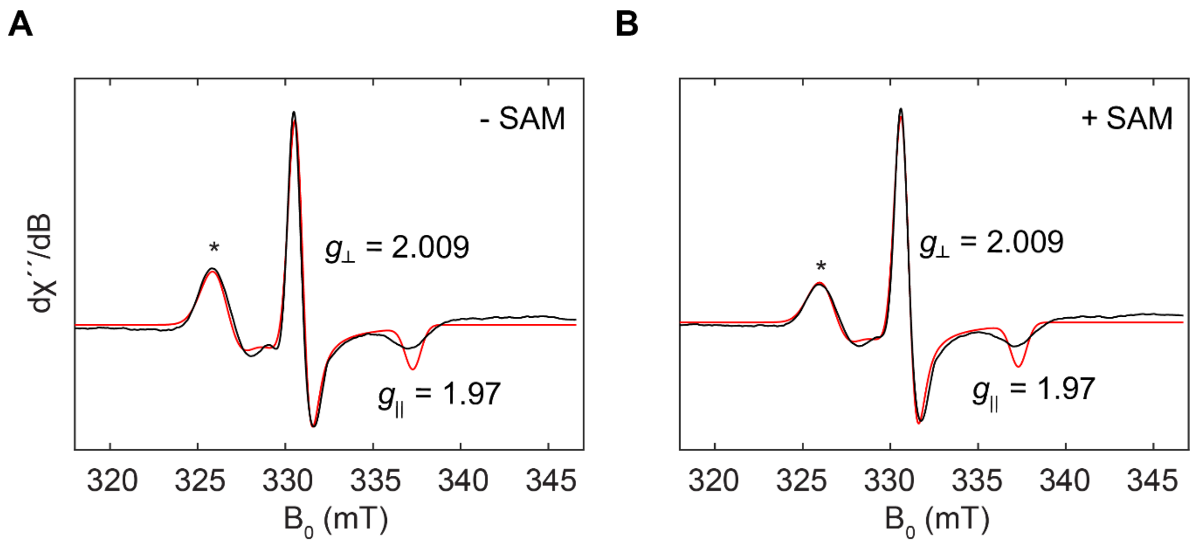

**Figure S9.** X-band EPR spectra of reduced PFL-AE. PFL-AE in the (**A**) presence or (**B**) absence of SAM following inhibition by NO. PFL-AE (50 μM) was reduced with 5 mM NaDT for 15 min, then mixed with 500 μM DEA-NONOate and incubated for 5 min before freezing in liquid $N_2$-cooled isopentane in an EPR tube. Black traces show experimental data and red traces the simulation. The asterisk denotes a small amount of signal corresponding to non-reduced PFL-AE DNIC. EPR conditions: microwave frequency, 9.3 GHz; modulation amplitude, 2 G; power 2 mW; temperature, 100 K. 30 scans were averaged for each spectrum.

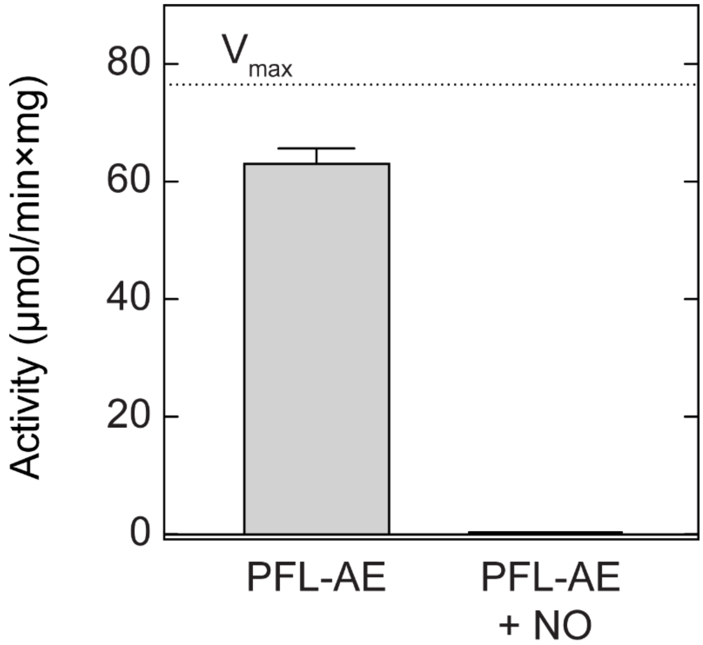

**Figure S10.** Inhibition of PFL-AE activity by NO. 20 μM PFL-AE was incubated with 100 μM NONOate for 30 minutes, then buffer exchanged by desalting. The treated PFL-AE was then used to activate PFL. PFL activity in the activation assay was monitored using the enzymatic coupled assay described in the main text. A control sample, consisting of untreated PFL-AE, was processed in the same way but without the addition of NONOate.

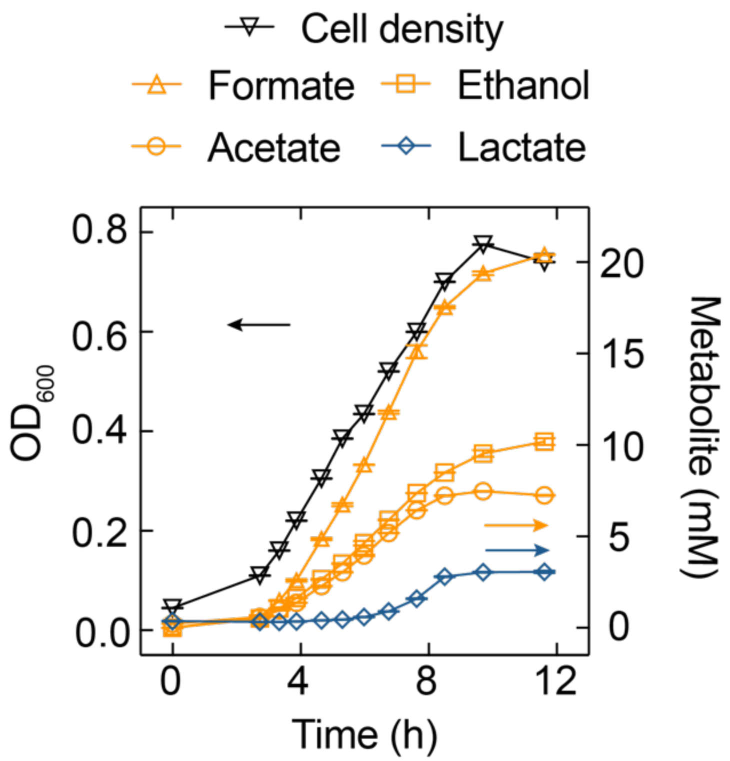

**Figure S11.** Metabolic analysis of anaerobically growing *E. coli*. Anaerobic cultures of *E. coli* in anaerobic conditions were monitored. Cell density was estimated from the $OD_{600}$ (black triangles) and extracellular concentration of the fermentation products lactate (blue diamonds), acetate (orange circles), ethanol (orange squares), and formate (orange triangles) were determined using enzymatic coupled assays. Error bars represent the span between two biological replicates.

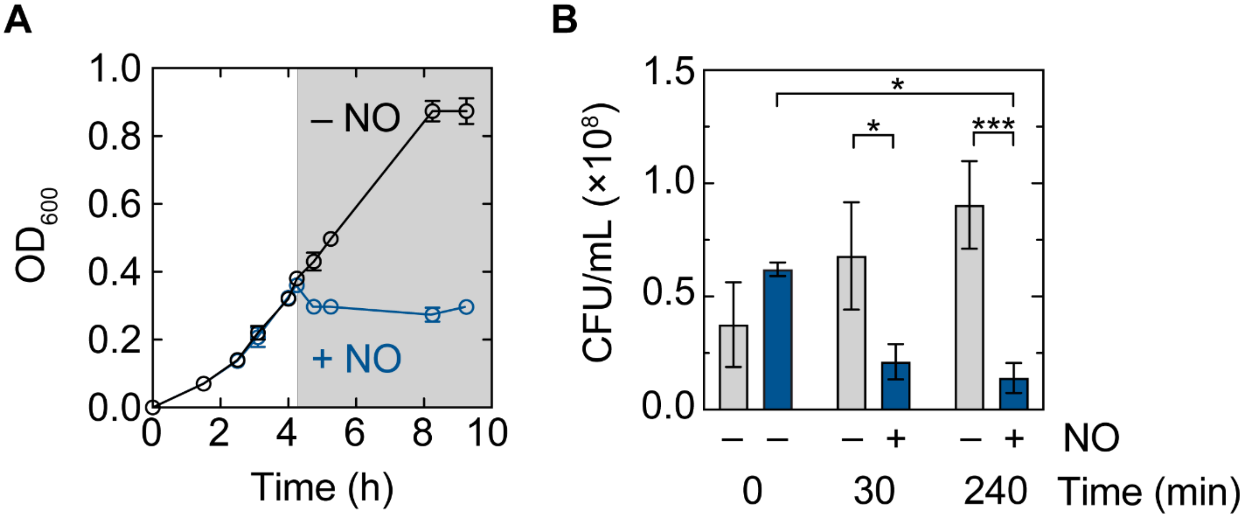

**Figure S12.** Bactericidal effects of NO in anaerobically growing *E. coli*. **A** Anaerobic cultures of *E. coli* were treated with 100 μM NONOate at mid-exponential phase (gray shaded area). Cell density was estimated from the $OD_{600}$ (black and blue circles). Error bars represent the standard deviation between three biological replicates. **B** Cell viability (CFU/mL) of anaerobically growing *E. coli* shown in panel **A**. Samples were taken at $OD_{600}$ 0.25 (0 min, pretreatment), 30 min and 240 min post 100 μM NONOate treatment. Error bars represent the standard deviation between three biological replicates. Comparative statical analysis was performed by two-way analysis of variance (ANOVA, * $p < 0.05$; *** $p < 0.001$).

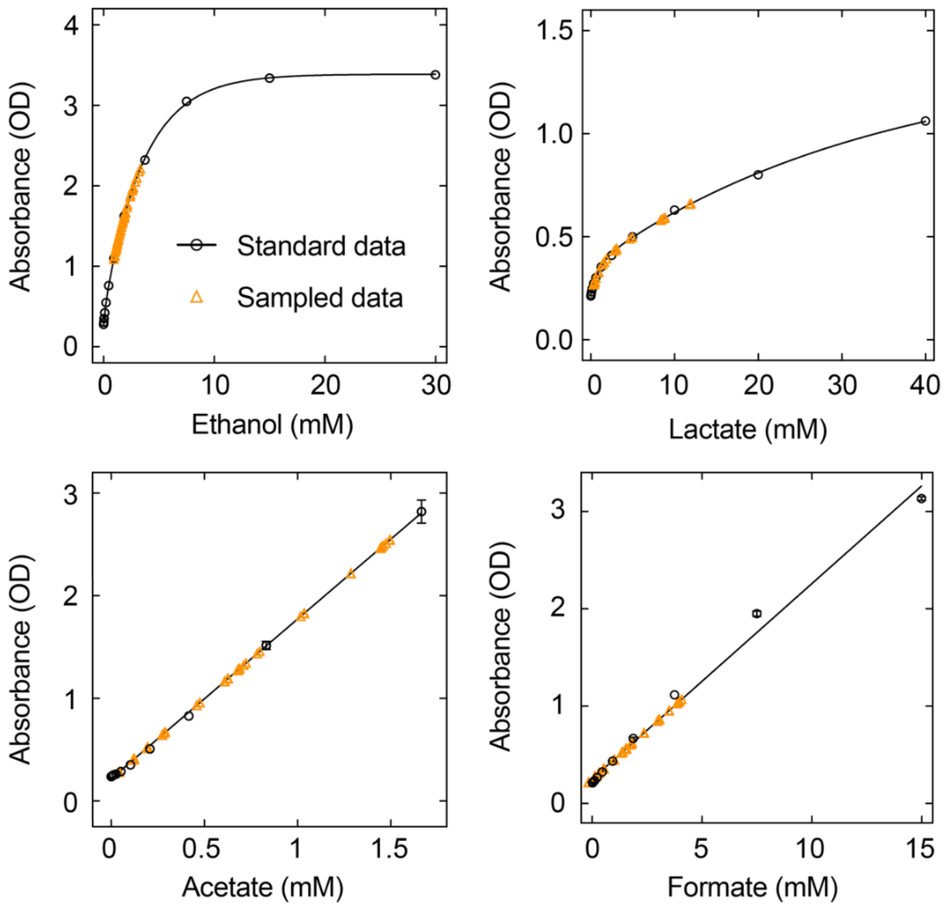

**Figure S13.** Calibration curves of *E. coli* fermentation products by enzyme coupled assays. Enzyme-coupled quantitation of ethanol (upper left), lactate (upper right), acetate (lower left), and formate (lower right) were fitted to calibration curves described in the Supplemental Methods. Assays (black circles) and corresponding experimental concentrations (orange triangles) are shown. The calibration curves were modeled (black lines) with either a linear regression (acetate, formate) or a two-phase association model (ethanol, lactate) where substrate inhibition has been reported. Error bars represent the span between two technical replicates. Where no error bars are reported, error was not resolved. Sampled data correspond to discreet values fitted into the calibration curve with no error bars.

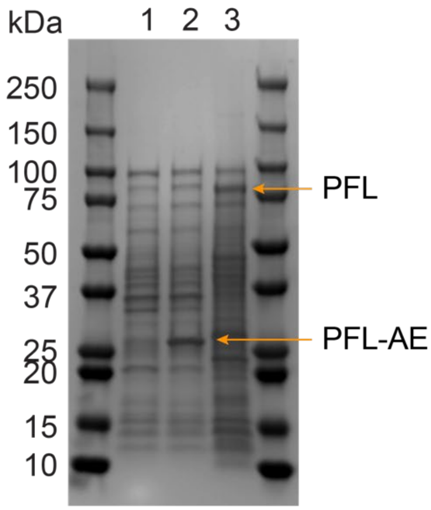

**Figure S14.** Over-expression of PFL and PFL-AE in anaerobically growing *E. coli* cells following induction with IPTG. Samples of anaerobic cultures of non-transformed *E. coli* (lane 1), *E. coli* over-expressing PFL-AE (lane 2) or PFL (lane 3) were resolved by SDS-PAGE. A molecular weight ladder is included for comparison.

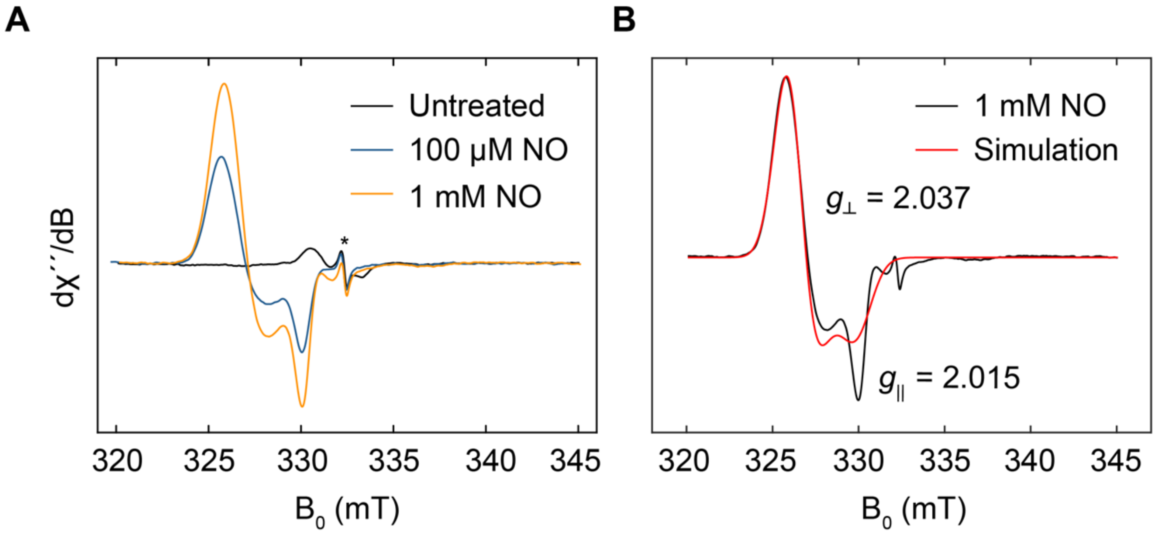

**Figure S15.** Whole-cell X-band EPR spectra of anaerobic E. coli cultures over-expressing PFL and treated with NONOate. **A** *E. coli* cultures were treated with 100 µM (blue), 1 mM, (orange) and without NONOate (black) for 10 min during stationary phase. Asterisk indicates a cavity artifact. **B** EPR spectra of *E. coli* cultures reacted with 1 mM NONOate from A (black) and simulation (red). EPR conditions: microwave frequency, 9.3 GHz; modulation amplitude, 2 G; power, 20 µW; temperature, 100 K. 30 Scans were averaged for each spectrum. * Depicts a background signal in the EPR cavity.

**Table S3. EPR simulation parameters for whole cells treated with NO.**

| Sample | $g_{\perp}$ | $g_{\parallel}$ | Fraction reduced, % |
|---|---|---|---|
| PFL + 1 mM NO | 2.03759(1) | 2.014(1) | 0 % |
| PFL + 100 μM NO | 2.03859(4) | 2.015(1) | 0 % |
| PFL-AE 100 μM NO | 2.03738(4) | 2.015(1) | 0 % |
| PFL-AE 1 mM NO | 2.03703(2) | 2.0144(7) | 25 % |
| | 2.00918(3) | 1.9709(1) | |
| Control + 1 mM NO | 2.03771(3) | 2.0144(7) | 14 % |
| | 2.00924(5) | 1.9749(1) | |
| Control + 100 μM NO | 2.03768(5) | 2.015(1) | 0 % |

Errors, reported in parenthesis for the last significant digit, are the standard deviations of the EasySpin simulation parameters.

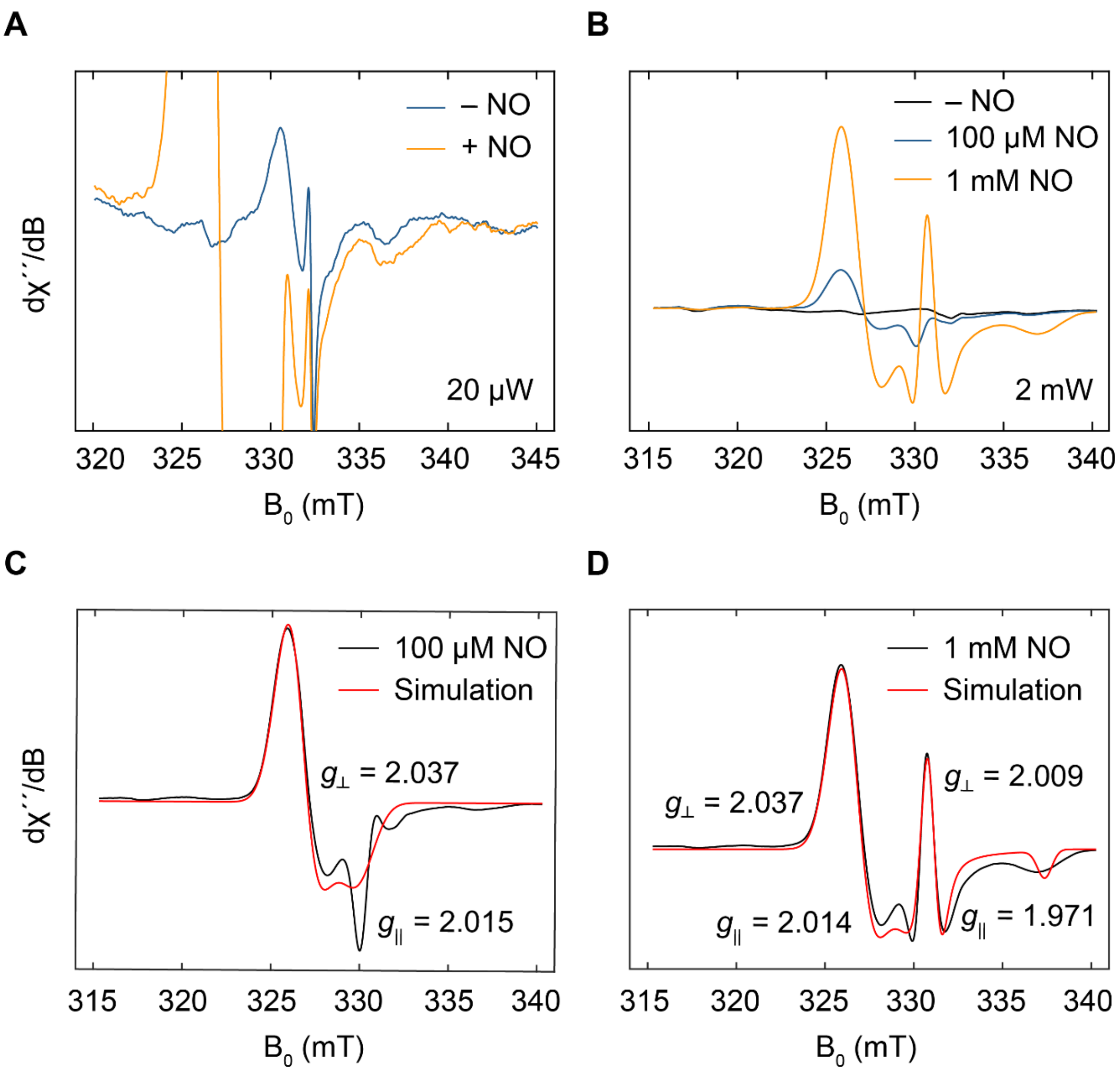

**Figure S16.** Whole-cell X-band EPR spectra of anaerobic E. coli cultures over-expressing PFL-AE with and without treatment with NONOate. **A** EPR spectra of *E. coli* cultures over-expressing PFL-AE from the same experiment described in **Figure 5B** in the main text, but recorded at 20 μW power, showing a close up of the glycyl radical. **B** EPR spectra of *E. coli* cultures over-expressing PFL-AE from the same experiment described in **Figure 5B**, a sample treated in the same way but with 1 mM NO is also examined. **C** EPR spectra of *E. coli* cultures over-expressing PFL-AE treated with 100 μM NO (black) and simulation (red) and **D** EPR spectra of *E. coli* cultures over-expressing PFL-AE treated with 1 mM NO (black) and simulation (red), corresponding to the same experiments shown on B. EPR conditions: microwave frequency, 9.3 GHz; modulation amplitude, 2 G; power, 20 μW (**A**) or 2 mW (**B**); temperature, 100 K. 30 scans were averaged for each sample.

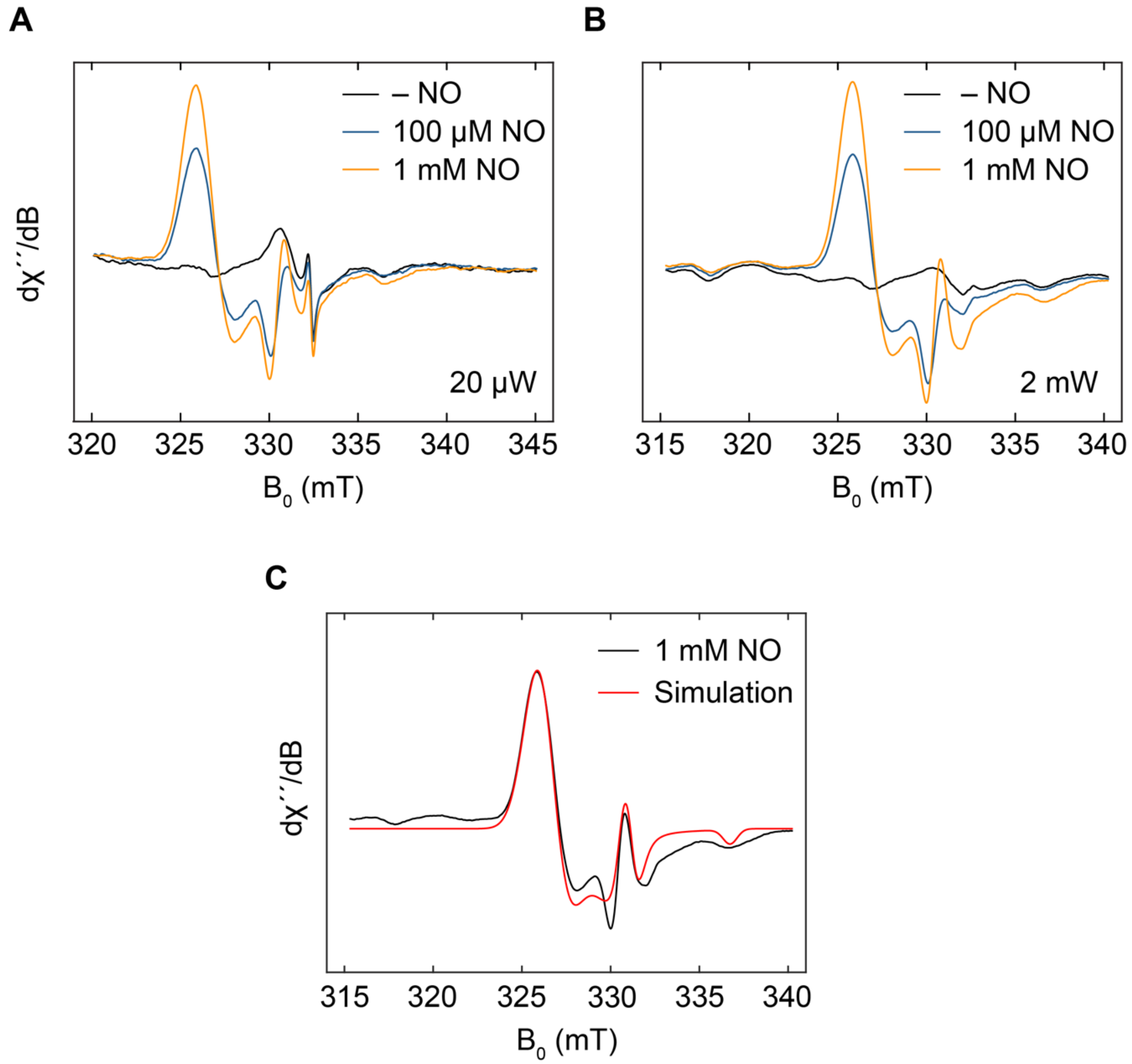

**Figure S17.** Whole-cell X-band EPR spectra of anaerobic E. coli cultures treated with NO. **A** EPR spectra of *E. coli* anaerobic cultures from the same experiment described in the main text **Figure 5B**, including non-treated samples and sample treated with 100 μM or 1 mM NONOate recorded at 20 μW or **B** recorded at 2 mW. **C** EPR spectra of *E. coli* cultures reacted with 1 mM NONOate from B (black) and simulation (red). EPR conditions: microwave frequency, 9.3 GHz; modulation amplitude, 2G; temperature, 100 K. 30 scans were averaged for each spectrum.

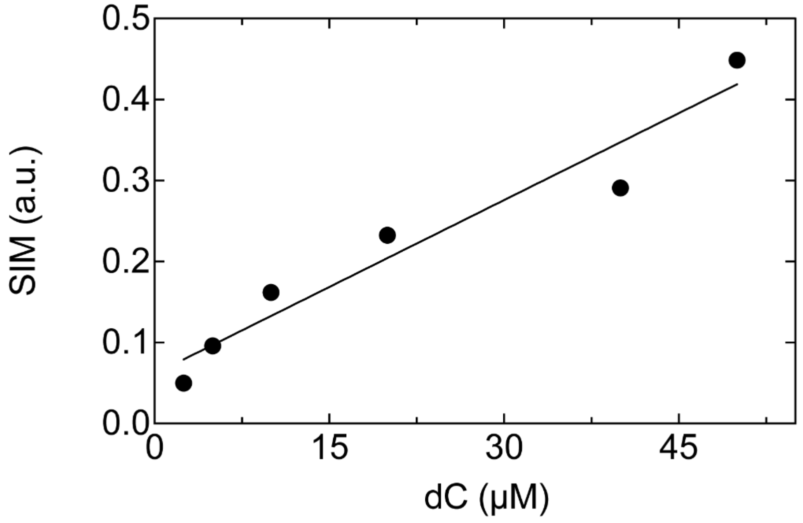

**Figure S18.** Deoxycytidine (dC) LC-MS calibration curve. Single ion monitoring (SIM) of dC (m/z 228.1 ± 50 ppm) peak integration versus dC concentration normalized to a 12.5 μM A (m/z 268.1 ± 50 ppm) internal standard to account for variations in injection volumes.

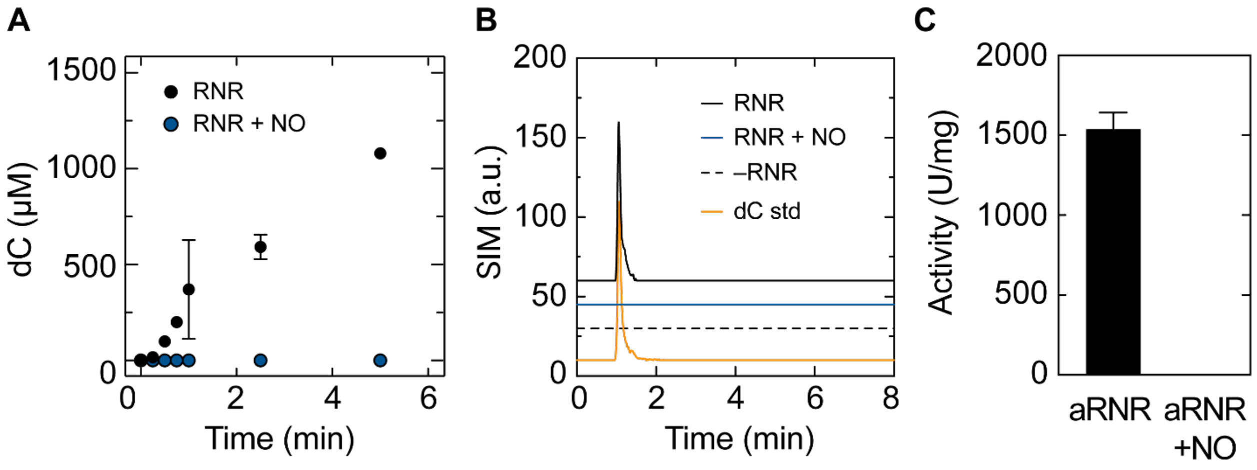

**Figure S19.** NO inactivation of *E. coli* RNR class III activity monitored by LC-MS. **A** Integrated SIM of dC over time, normalized to A for aRNR (black) and aRNR treated with 500 µM NONOate for 10 min (blue). **B** LC-MS traces of SIM channels of 30 µM dC standard (orange), assay buffer without aRNR (black dashed), and 5 min reaction time point for aRNR (black) or aRNR treated with 500µM NONOate for 10 min (blue). Offsets added for comparison and clarity. **C** Specific activity of aRNR vs aRNR incubated with 500µM NO for 10 minutes. One unit (U) of activity is defined as 1 nmol/min dC. The detection limit for activity is > 3.5 U/mg.

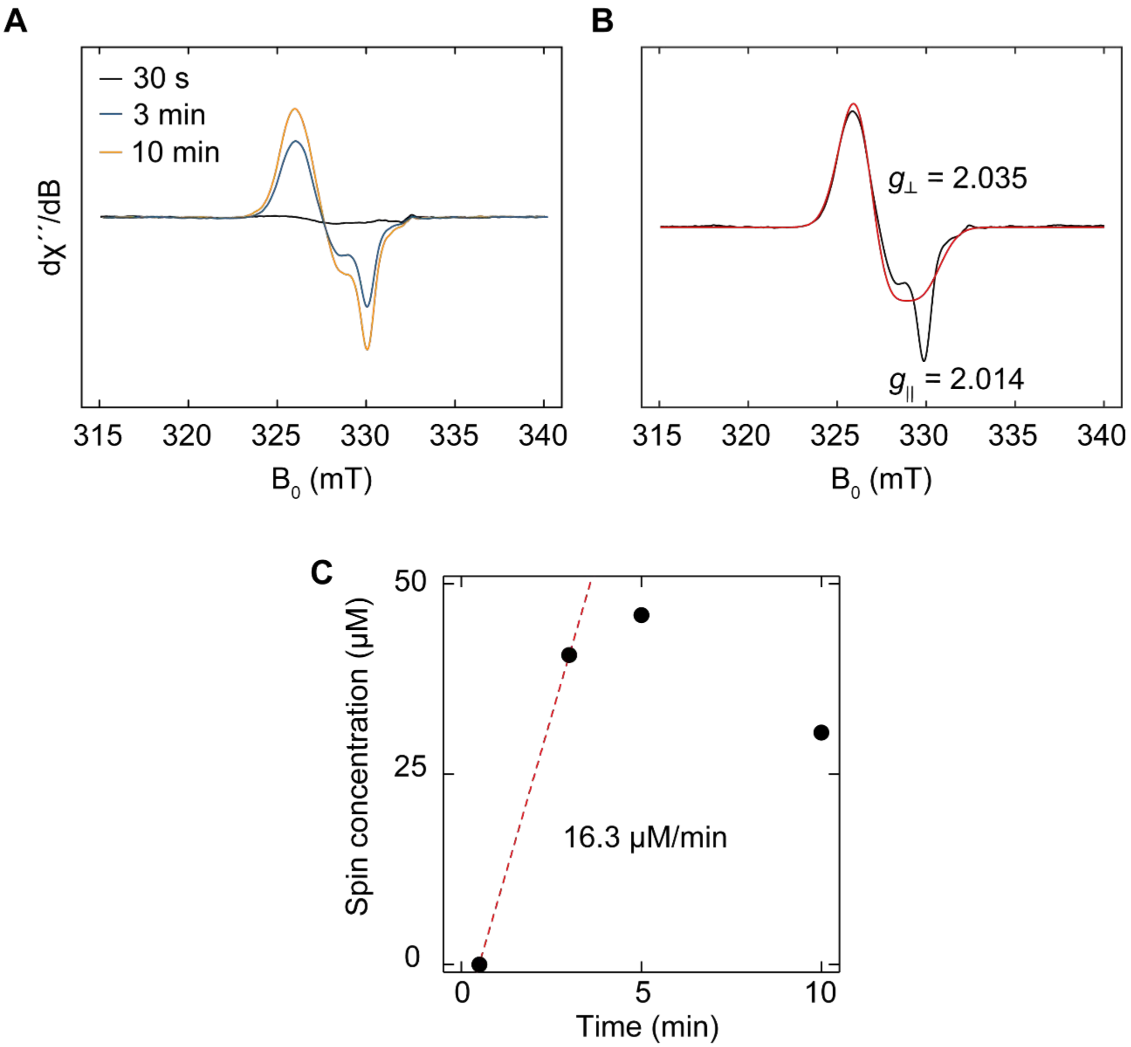

**Figure S20.** X-band EPR spectra of RNR-AE reacted with NO. **A** EPR spectra of 50 µM RNR-AE mixed with 500 µM NONOate and incubated for 30 s (black), 3 min (blue), and 10 min (orange). **B** EPR spectra of RNR-AE reacted with NONOate for 10 minutes from panel A (black) and simulation (red). EPR conditions: microwave frequency, 9.3 GHz; modulation amplitude, 2 G; power, 2 mW; temperature, 100 K. 30 scans were averaged for each spectrum. **C** Kinetics of RNR-AE DNIC formation. Spin concentrations were computed from the double integral of the first harmonic signal of the EPR spectra shown in panel A and referenced to a 4-hydroxy-TEMPO standard.

**Table S4. EPR simulation parameters for RNR-AE treated with NO.**

| Sample | $g_{\perp}$ | $g_{/\!/}$ | Concentration (μM) | Spin/protein (%) |
|---|---|---|---|---|
| RNR-AE 10 min | 2.00355 (1) | 2.014 (2) | 30.5 | 61 |
| 5 min | | | 45.9 | 92 |
| 3 min | | | 40.7 | 81 |
| 30 s | | | 0 | 0 |
| RNR-AE + SAM | 2.0356(1) | 2.014(2) | 3.4 | 7 |
| Sample | $g_{\perp}$ | $g_{/\!/}$ | Concentration (μM) | Fraction reduced (%) |
| rRNR-AE | 2.0100 (1) | 1.9710 (1) | 5.0 | 64 % |
| rRNR-AE + SAM | 2.00988 (2) | 1.9706 (2) | 4.8 | 66 % |

Errors, reported in parenthesis for the last significant digit, are the estimated uncertainty of the EasySpin non-linear least squares fit simulation parameters.

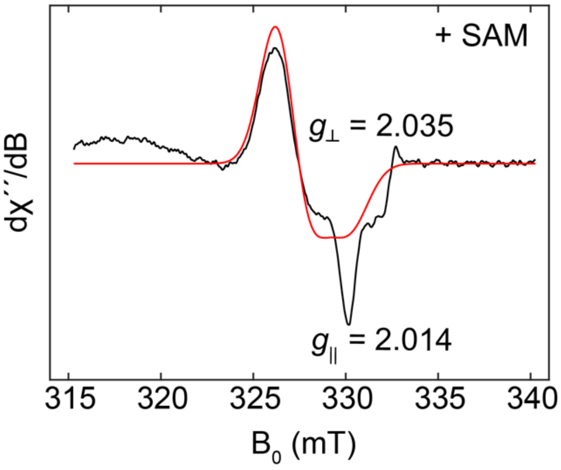

**Figure S21.** X-band EPR spectrum of RNR-AE reacted with NONOate in the presence of SAM. 50 μM RNR-AE was reacted with 500 μM NONOate over 5 min in the presence of SAM then frozen in liquid $N_2$-cooled isopentane in an EPR tube. Black trace shows experimental data and red trace simulated data. EPR conditions: microwave frequency, 9.3 GHz; modulation amplitude, 2 G; power 2 mW; temperature, 100 K. 30 scans were averaged.

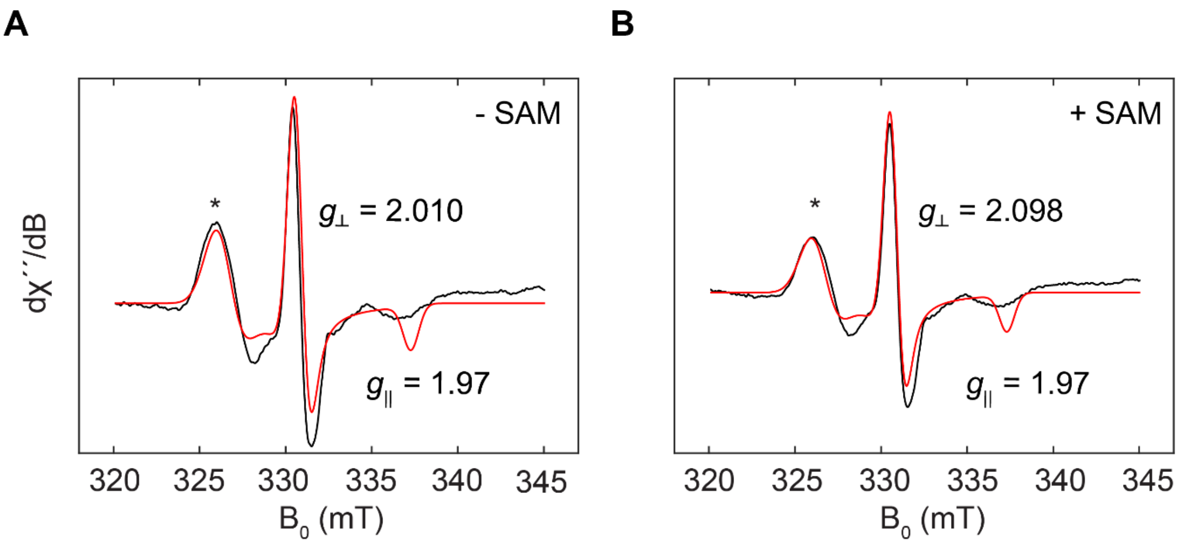

**Figure S22.** X-band EPR spectra of sodium dithionite (NaDT) reduced RNR-AE with NO. X-band EPR spectra of RNR-AE 50 μM RNR-AE reduced with 5 mM sodium dithionite (NaDT) for 10 min and then reacted with 500 μM DEA-NONOate for 5 min with (orange) or without (black) 2 mM SAM. EPR conditions: microwave frequency, 9.3 GHz; modulation amplitude, 2 G; power, 2 mW; temperature, 100 K. 30 scans were averaged for each spectrum.

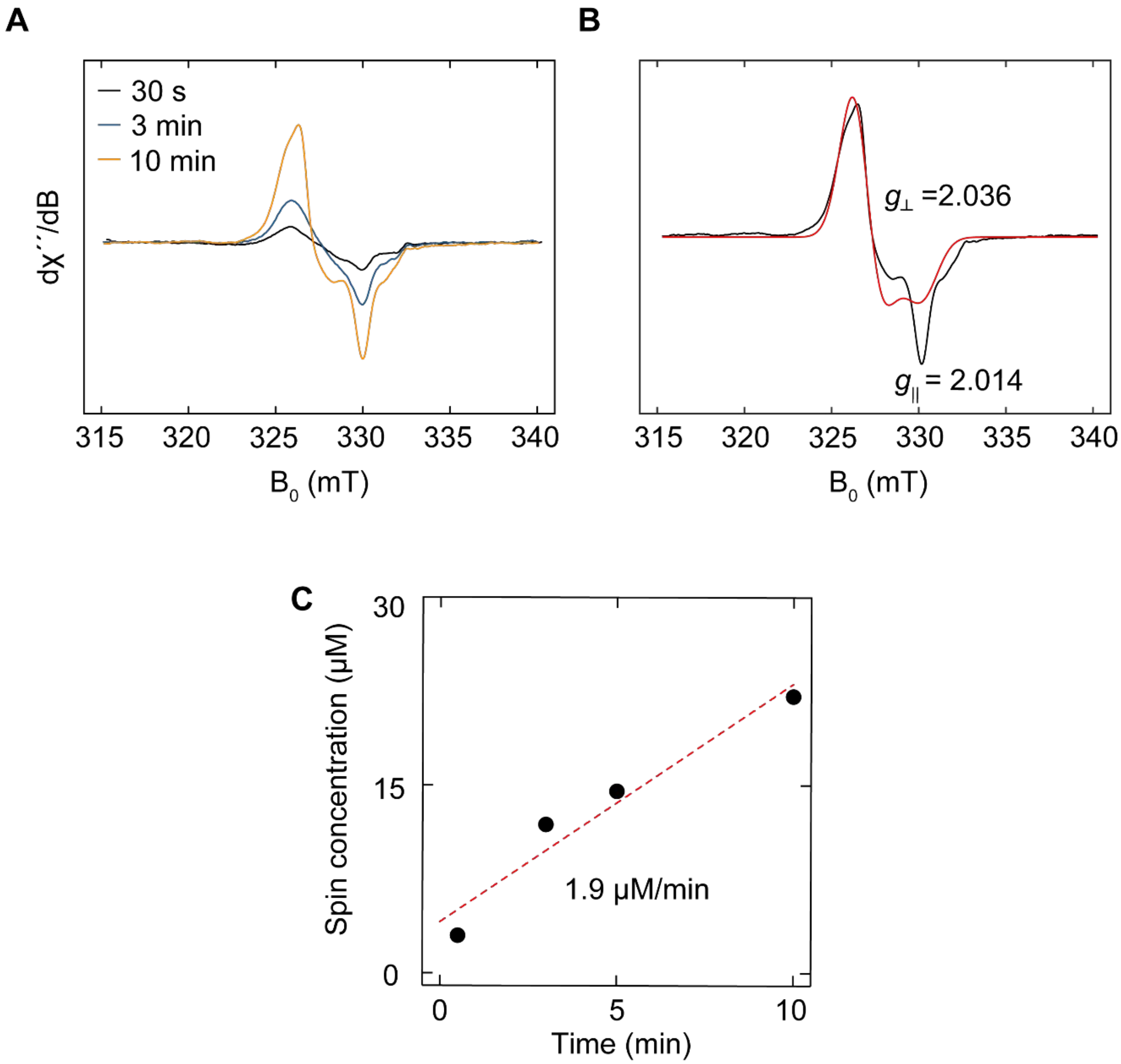

**Figure S23.** X-band EPR spectra of NikJ reacted with NO. **A** EPR spectra of 50 μM NikJ mixed with 500 μM NONOate for 30 s (black), 3 min (blue), and 10 min (orange). **B** EPR spectra of NikJ reacted with NONOate for 10 minutes from A (black) and simulation (red). EPR conditions: microwave frequency, 9.3 GHz; modulation amplitude, 2 G; power, 2 mW; temperature, 100 K. 30 scans were averaged for each spectrum. **C** Kinetics of NikJ DNIC formation, spin concentrations were computed from the double integral of the first harmonic signal of the EPR spectra shown in A and referenced to a 4-hydroxy-TEMPO standard.

**Table S5. EPR simulation parameters for NikJ treated with NO.**

| Sample | $g_{\perp}$ | $g_{/\!/}$ | Concentration (μM) | Spin/protein (%) |
|---|---|---|---|---|
| NikJ 10 min | 2.00361 (1) | 2.0142 (9) | 22.1 | 44 |
| 5 min | | | 14.5 | 29 |
| 3 min | | | 11.9 | 24 |
| 30 s | | | 3 | 6 |
| NikJ + SAM | 2.0364(1) | 2.014(1) | 17.9 | 36 |
| Sample | $g_{\perp}$ | $g_{/\!/}$ | Concentration (μM) | Fraction reduced (%) |
| rNikJ | 2.00925 (4) | 1.9710 (1) | 24.8 | 16 |
| rNikJ + SAM | 2.00957 (2) | 1.9700 (1) | 22.2 | 81 |

Errors, reported in parenthesis for the last significant digit, are the standard deviations of the EasySpin simulation parameters.

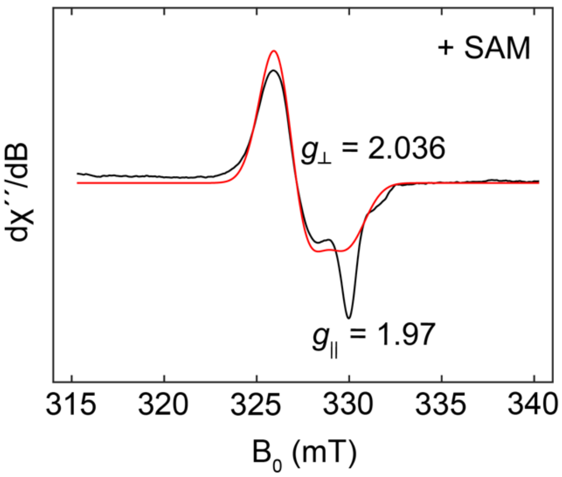

**Figure S24.** X-band EPR spectrum of NikJ reacted with NONOate in the presence of SAM. 50 μM NikJ was reacted with 500 μM NONOate over 5 min in the presence of SAM then frozen in liquid $N_2$-cooled isopentane in an EPR tube. Black trace shows experimental data and red trace simulated data. EPR conditions: microwave frequency, 9.3 GHz; modulation amplitude, 2 G; power 2 mW; temperature, 100 K. 30 scans were averaged.

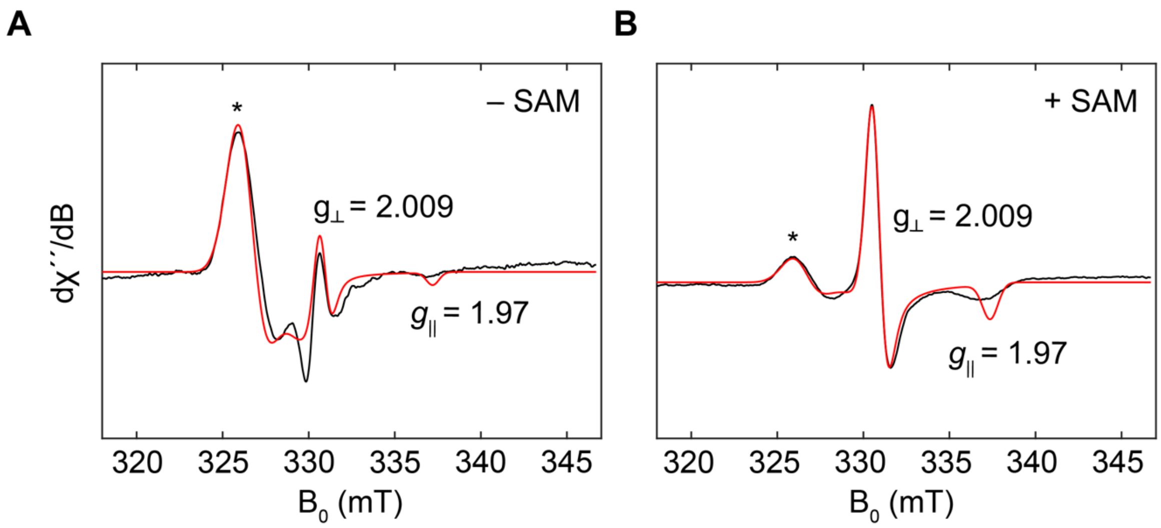

**Figure S25.** X-band EPR spectra of sodium dithionite (NaDT) reduced NikJ with NO. EPR spectra of 50 µM NikJ reduced with 5 mM NaDT over 15 min with (**A**) or without 2 mM SAM (**B**) then mixed with 500 µM DEA-NONOate and incubated for 5 min. Black trace shows experimental data and red trace simulated data. EPR conditions: microwave frequency, 9.3 GHz; modulation amplitude, 2 G; power, 2 mW; temperature, 100 K. 30 scans were averaged for each spectrum.

### References

1. Hewitt, C. O.; Eszes, C. M.; Sessions, R. B.; Moreton, K. M.; Dafforn, T. R.; Takei, J.; Dempsey, C. E.; Clarke, A. R.; Holbrook, J. J. A General Method for Relieving Substrate Inhibition in Lactate Dehydrogenases. *Prot. Eng. Des. Sel.* **1999**, *12*, 491–496, DOI: 10.1093/protein/12.6.491.

2. Shore, J. D.; Theorell, H. Substrate Inhibition Effects in the Liver Alcohol Dehydrogenase Reaction. *Arch. Biochem. Biophys.* **1966**, *117*, 375–380, DOI: 10.1016/0003-9861(66)90425-5.

3. Cáceres, J. C.; Dolmatch, A.; Greene, B. L. The Mechanism of Inhibition of Pyruvate Formate Lyase by Methacrylate. *J. Am. Chem. Soc.* **2023**, *145*, 22504–22515, DOI: 10.1021/jacs.3c07256.

4. Knappe, J.; Volker Wagner, A. F. Glycyl Free Radical in Pyruvate Formate-Lyase: Synthesis, Structure Characteristics, and Involvement in Catalysis. In *Methods in Enzymology*; Elsevier, 1995; Vol. 258, pp 343–362, DOI: 10.1016/0076-6879(95)58055-7.
